# High-quality genome assemblies of diploid *Bromus* species enhance understanding of genome complexity and uncover large DNA satellite structures

**DOI:** 10.64898/2026.01.08.698408

**Authors:** Raju Chaudhary, Yuanyuan Ji, Kevin C. Koh, Sampath Perumal, Zhengping Wang, Irfan Iqbal, Venkatesh Bandi, Bill Biligetu, Andrew G. Sharpe

## Abstract

Genus *Bromus* includes important cool-season forage grasses, but large genome sizes and complex ploidy hinder genomic studies. Here we carried out long read sequencing of two diploid genomes, *Bromus riparius* and *Bromus squarrosus*, generating highly contiguous assemblies with contig N50 values of 294.76 Mb and 98.78 Mb, respectively. These assemblies uncovered high-order centromeric repeats and two unique large sub-telomeric repeats on chromosome 5 and 7. Repeat expansion in *Bromus* appears relatively recent compared to speciation events (∼9.75 million years ago); yet strong synteny persists across species. Transposable elements such as Angela, SIRE, Athila, CRM, and Retand were identified as major contributors to genome expansion in related *Bromus* species. Population structure analysis resolved six major subpopulations, with wild relatives exhibiting a higher genetic divergence than cultivated types. Hybrid bromegrass lines showed a higher proportion of genomic contribution from *B. inermis* than *B. riparius.* Together, these genomic resources provide a foundation for investigating traits of interest and advancing bromegrass breeding.

## Main

Genus *Bromus* (family Poaceae) comprises over 400 annuals, biennials and perennial grasses (Leighton and Harms, 2014). Cultivated *Bromus* species are believed to have originated from Eurasia (Brouillet et al., 2010). Taxonomic classification of *Bromus* has been complex, with multiple classifications proposed by different taxonomists. Among these, the seven-section classification by Smith (1970) has been widely adopted due to the extensive detail of its structure. The *Pnigma* Dumort. is the major section comprising of more than 60 species, followed by the *Bromus* section containing 40 species, and other sections being *Ceratochloa* (P. Beauv.) Griseb., *Genea* Dumort., *Neobromus* (Shear.) Hitchcock, *Nevskiella* (Krecz &Vved.) and *Boissiera* (Hochst. Ex Steudel). These sections exhibit significant evolutionary divergence, particularly in genome size and ploidy (Armstrong, 1991). Section *Pnigma* is an agriculturally important section due to the presence of species with excellent nutritional value and yield potential (Casler et al., 2000). This section includes *B. inermis* Leyss., *B. riparius* Rehm. and *B. erectus* Huds. Among these, *B. inermis* (smooth bromegrass) and *B. riparius* (meadow bromegrass), which were introduced in North America around 1888 and 1960 respectively (Anstey, 1986), are extensively cultivated as forage crops in western Canada (Biswas et al., 2020). *Bromus inermis* is valued for high hay yield, whereas meadow bromegrass is preferred for its rapid regrowth. Interspecific hybrids between these species have been developed to combine these desirable traits, enhancing forage production and adaptability (Coulman, 2004).

Most of the wild species found in North America are either diploid or tetraploid whereas *Bromus* species found in Eurasia are of higher ploidy (Armstrong, 1991). A limited assessment of genetic variation have been explored among the smooth bromegrass and meadow bromegrasses using Random Amplified Polymorphic DNA (RAPD) and Amplified Fragment Length Polymorphism (AFLP) markers (Ainouche et al., 1999; Diaby and Casler, 2003; Ferdinandez and Coulman, 2002; Li et al., 2006) as well as a Genotyping-by-Sequencing (GBS) approach (Lawrence et al., 2017). These studies revealed moderate to high genetic diversity, highlighting the potential for increasing genetic diversity and subsequently improving bromegrass breeding programs. Recently, genomes of annual bromegrass such as *B. tectorum* (Revolinski et al., 2023) and *B. sterilis* (Christenhusz et al., 2024) have been published to aid the characterization of *Bromus* species; however, no information is available on the cultivated bromegrass species.

Cytological studies have further highlighted the complexity of *Bromus* genomes. Satellite chromosomes were reported in *B*. *inermis* (Ghosh and Knowles, 1964; Tuna et al., 2004), and similar satellites were identified in diploid perennial such as *B. erectus* and *B. variegatus* (Tuna et al., 2006), as well as tetraploid annual *B. hordeaceus* (Ainouche *et al*., 1999). Chromosome pairing analysis and karyotype studies suggested presence of multiple subgenomes in higher species of genus *Bromus*; however, comprehensive subgenome assignment remain incomplete (Armstrong, 1991; Tuna *et al*., 2004). Variation in chromosome length and genome size (Tuna *et al*., 2006) further complicates species classification, especially in cultivated lines where hybridization and variation in ploidy levels are common. It is speculated that the *Bromus* genome consists of diploid to duodecaploid species, with octaploid being the common among the cultivated types (Williams et al., 2011).

The limited understanding on the genomic organization in *Bromus* (Armstrong, 1991), unavailability of reference genome for cultivated bromegrass, and lack of precise genetic relationship information within each cultivated bromegrass is limiting bromegrass breeding and genetic improvement. In addition, the observed variabilities in genomic content, chromosome length, and ploidy forms within the genus further provides a valuable yet underexplored opportunity for studying underlying genome biology (Williams *et al*., 2011). Harnessing this variation could significantly contribute to the improvement of agriculturally important bromegrasses. Here, we sequenced two diploid bromegrass accessions representing different Bromus sections, *Bromus riparius* (PI440215) and *Bromus squarrosus* (PGR7039), providing high-quality genome assemblies with large satellite chromosome structures identified, and established the relationships among *Bromus* species.

## Results

### Genomes assemblies of diploid bromegrass accessions

Two diploid genomes of *Bromus*, representing *B. riparius* (accession PI440215) and a weedy relative *B. squarrosus* (accession PGR7039), were sequenced using PacBio long-read sequencing technology. Both genomes contain n = 7 chromosomes, however the genome size is remarkably different between the species (**Supplementary Fig. 1 & 2**) with 2,407.6 Mb and 5,104.7 Mb for PI440215 and PGR7039 respectively (Marçais and Kingsford, 2011; Vurture et al., 2017). The High-Fidelity (HiFi) reads with a coverage of 70× for PI440215 and 32× for PGR7039 were assembled using hifiasm v.0.19.5 (Cheng et al., 2022) and the base assemblies were scaffolded to a chromosome level using Hi-C technology (Burton et al., 2013). The size of the primary assembly for PI440215 was 2,509.35 Mb with a contig N50 of 294.76 Mb (**Table 1, Supplementary Table 1**) which was further scaffolded to a chromosome level with an N50 of 350.02 Mb. The size of the primary assembly for PGR7039 was 4,362.82 Mb with a contig N50 of 98.78 Mb, which further scaffolded into 7 chromosomes with an N50 of 641.9 Mb (**Fig. 1**). The scaffolding was able to anchor more than 95.77% and 97.91% of genomes to a chromosome level for PI440215 and PGR7039 respectively. The Benchmarking Universal Single-Copy Orthologs (BUSCO) analysis (Simão et al., 2015) using embryophya_odb10 database suggested more than 98.14% of the BUSCO genes are completely present in these genomes (**Table 1**). Further, Illumina based short reads (∼79× for PI440215 and 27× for PGR7039) were mapped to these genomes which showed mapping percentage of 99.69% and 99.18% in PI440215 and PGR7039 respectively (**Supplementary Table 2**). HiFi read mapping further demonstrated uniform coverage across genome, underscoring the high quality of both assemblies (**Fig. 1**).

**Table 1.**
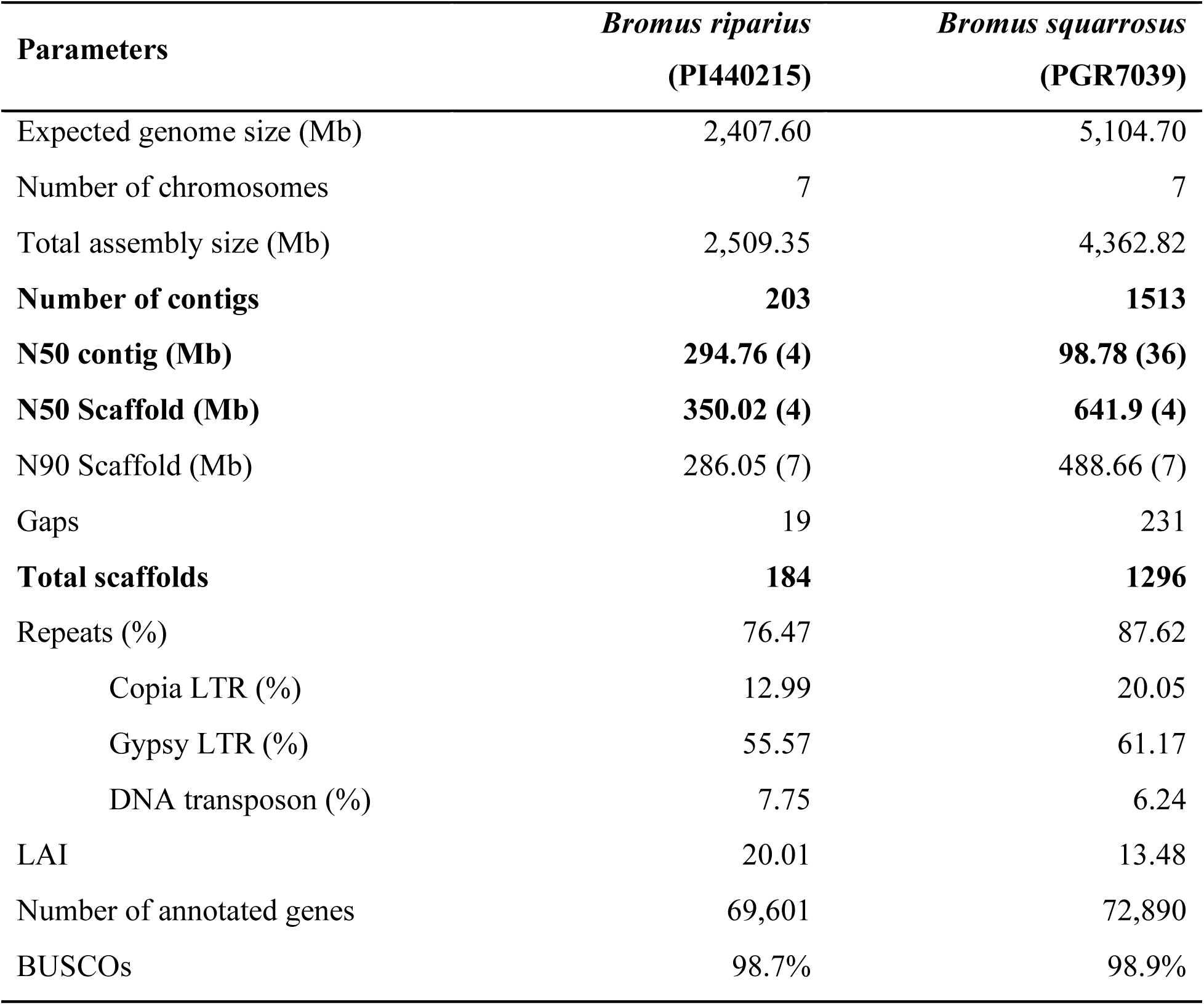
Summary statistics of the genome assemblies of diploid bromegrass accessions.

| **Parameters** | ***Bromus riparius***<br>**(PI440215)** | ***Bromus squarrosus***<br>**(PGR7039)** |
| --- | --- | --- |
| Expected genome size (Mb) | 2,407.60 | 5,104.70 |
| Number of chromosomes | 7 | 7 |
| Total assembly size (Mb) | 2,509.35 | 4,362.82 |
| **Number of contigs** | **203** | **1513** |
| **N50 contig (Mb)** | **294.76 (4)** | **98.78 (36)** |
| **N50 Scaffold (Mb)** | **350.02 (4)** | **641.9 (4)** |
| N90 Scaffold (Mb) | 286.05 (7) | 488.66 (7) |
| Gaps | 19 | 231 |
| **Total scaffolds** | **184** | **1296** |
| Repeats (%) | 76.47 | 87.62 |
| Copia LTR (%) | 12.99 | 20.05 |
| Gypsy LTR (%) | 55.57 | 61.17 |
| DNA transposon (%) | 7.75 | 6.24 |
| LAI | 20.01 | 13.48 |
| Number of annotated genes | 69,601 | 72,890 |
| BUSCOs | 98.7% | 98.9% |

**Fig. 1:**
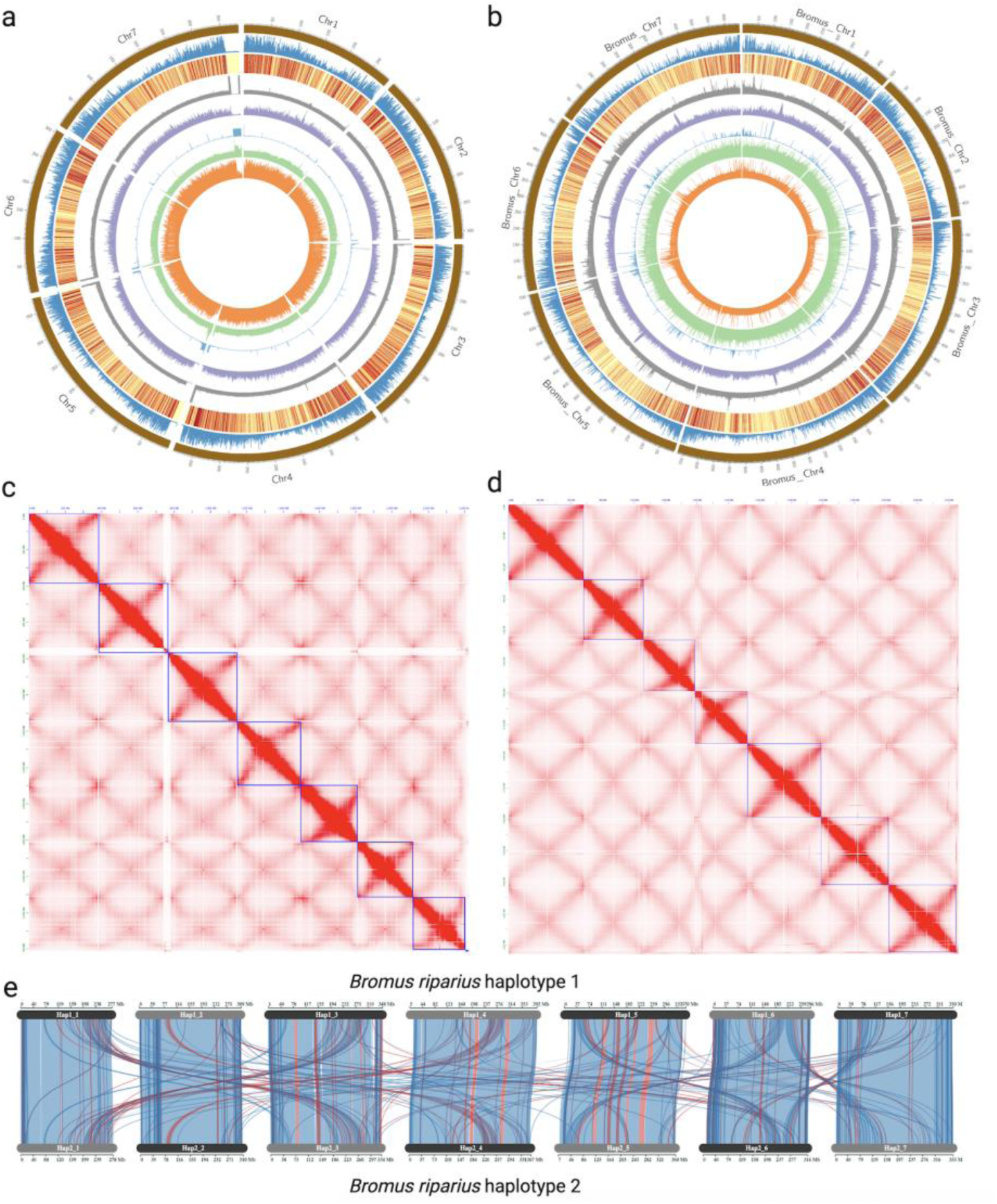
Genomes of diploid bromegrass species *Bromus riparius* (PI440215) and *B. squarrosus* (PGR7039). (a, b) Circos plots of PI440215 (a) and PGR7039 (b) showing seven chromosomes (outer track - brown), gene density (blue), gene expression as a heatmap (yellow to red), repeat distribution (grey), LTR distribution (purple), tandem repeat distribution (light blue), Illumina reads (green) and distribution of HiFi reads (orange). (c, d) Hi-C contact maps confirming scaffolding of primary assembly of PI440215 and PGR7039 respectively. e) Collinearity analysis between haplotype1 and haplotype 2 of PI440215, showing structural rearrangements including chromosomal inversions (red) and gene duplications originating from different chromosomes (blue).

The high coverage of HiFi reads enabled the construction of a haplotype-resolved assembly of *B. riparius* (**Fig. 1**). Both haplotypes are comparable in size, with haplotype 1 being slightly larger but less contiguous than haplotype 2 (**Supplementary Table 1**). HiC data were employed to scaffold the assemblies to produce a reference-quality phased genome assembly (**Supplementary Fig. 3**). The assembled genomes have BUSCO score of 98.2 and 96.7% in haplotype 1 and haplotype 2, respectively (**Supplementary Fig 4**). Comparative analysis of the two haplotypes suggested presence of several structural differences, including multiple inversions (**Fig. 1e**).

### Repeat dynamics in bromegrass genomes

The genomes were further analysed for the repeat contents using the EDTA pipeline (Ou et al., 2019) on each genome independently. The analysis predicted 76.47 % of the genome consists of repeat elements in PI440215 and 87.62 % in PGR7039 (**Supplementary Table 3**). Gypsy is the most abundant repeats, belonging to the Long Terminal Repeat (LTR) class, comprising 55.57% of the PI440215 and 61.17% of the PGR7039 genome. Copia elements accounts for 12.99% and 20.05% of the genomes PI440215 and PGR7039, respectively. Analysis of intact copy LTRs identified 40,076 full-length LTR elements in PI440215 and 52,601 in PGR7039. The LTR Assembly Index (LAI) (Ou et al., 2018), which is used to assess the quality of an assembly based on the resolution of repetitive LTRs, is also high with a value of 20.01 and 13.48 for PI440215 and PGR7039, respectively, suggesting reference quality of these genomes (**Table 1**). Among these LTR, Angela, SIRE, Athila, CRM, Retand and Tekay are the major repeat clades (**Supplementary Table 4)**. CRM (3.22 times), SIRE (2.38 times), Angela (4.18 times), Tork (12.56 times), and Galadriel (71 times) have very high copy number in PGR7039 compared to PI440215 (**Supplementary Fig. 5**). All the LTR repeats have proliferated recently in PI440215 compared to PGR7039, however, Angela and SIRE elements have evolved multiple times in PGR7039 (**Supplementary Fig. 6**). The distribution of centromere associated CRM elements are spread over a larger genomic region in PGR7039 (13.79 Mb - 22.13 Mb) in comparison to PI440215 5.3Mb - 9.2 Mb), which could suggest expansion of centromeric and peri-centromeric spaces in the PGR7039 (**Fig. 2**). A variability in tandem repeat exists across different centromeres, suggesting diversity in centromere specific tandem repeats (**Fig. 2a and Fig. 2b, Supplementary Table 5 & 6**). The canonical telomeric repeat TAGGGTT and TAGGGTTT were found in both PGR7039 and PI440215 genomes. In PI440215, telomeres were assembled on only three chromosomes, each at a single end, whereas in PGR7039, telomeres were assembled at both ends of all seven chromosomes (**Supplementary Fig. 7**). Notably, chromosomes 1 and 2 of PGR7039 exhibit a strong enrichment of telomeric repeats in its centromeric region. A total of 22,549 telomeric repeat copies were detected within a 6.1 Mb region on chromosome 1, while 51,374 telomeric repeats within a 0.59 Mb region on chromosome 2 at the proximal edge of the centromere (**Supplementary Table 6**) which may reflect remnants of an ancient chromosomal translocation or fusion. BLAST analysis using the *H. vulgare* 45S rDNA (HQ825319.1) (Georgiev and Karagyozov, 2012) identified multiple high-identity matches distributed across different chromosomes, with pronounced clustering on chromosome 1 in both PI440215 and PGR7039. This distribution pattern suggests presence of nucleolar organizer regions (NORs) in these genomes (**Supplementary Fig. 7**).

**Fig. 2:**
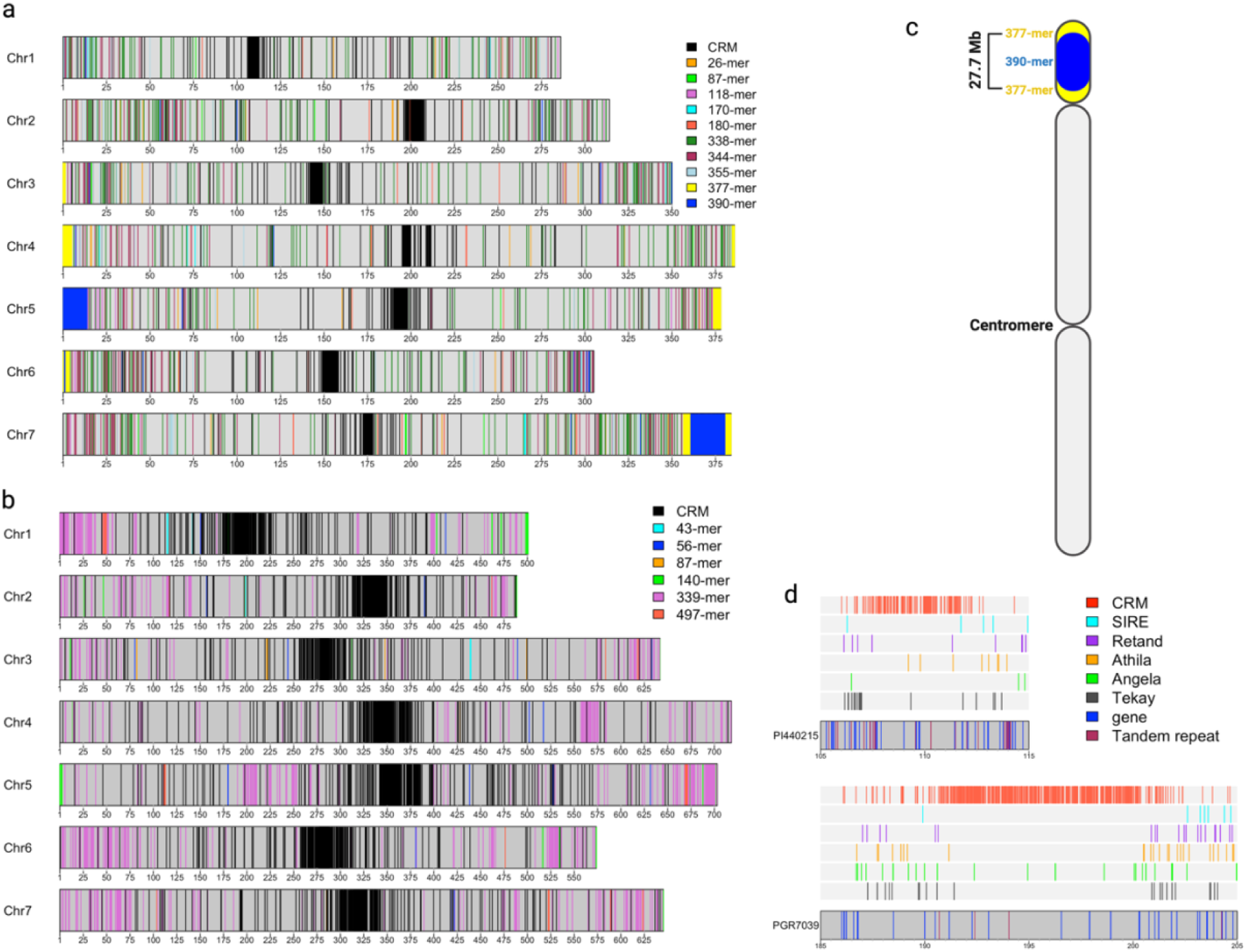
Distribution of tandem repeat elements in diploid genomes of bromegrass PI440215 and PGR7039. (a) Linear plots of the seven chromosomes of PI44015 showing distribution of tandem repeats with different monomer sizes. Blue blocks represent sub-telomeric repeats of a 390-mer, which are flanked by 377-mer repeats (yellow) on chromosome 7 of PI440215, while black blocks represent distribution of centromere specific repeats. (b) Linear plots of the seven chromosomes of PGR7039, showing the distribution of distinct tandem repeats, with black blocks marking centromere species repeats. (c) Schematic diagram of the satellite repeat structure in *Bromus riparius,* illustrating the organization of the 377-mer (yellow) surrounding the 390-mer (blue). (d) Comparison of full-length LTR retrotransposons and tandem repeats in the centromeric region of chromosome 1 in PI440215 and PGR7039, showing expansion of CRM retrotransposons contributing to increased centromere length.

In concordance with the previous cytological study (Tuna et al., 2001a) of PI440215, we were able to characterize sub-telomeric repeat (STR) structures on multiple chromosomes including two very large ones on chromosome 5 and chromosome 7, respectively. The HiC analysis confirmed their placement on both chromosomes (**Supplementary Fig. 8**). The STR has peculiar structure on chromosome 7 with repeats of length 390 bp flanked by another monomer repeat of 377 bp monomers (**Fig. 2c**). The size of the STR on chromosome 7 was 27.7 Mb and on chromosome 5 was 12.5 Mb with polymorphism among the tandem repeats. The frequency of such tandem repeats is very low in other studied *Bromus* species (**Supplementary Fig. 9**). This could suggest a unique feature to the PI440215 genome. In the case of PGR7039, a conservation of 339 bp tandem repeats were predicted around the telomeres for most of the chromosomes and several repeat monomers might represent satellite repeats different from the STRs in PI440215 (**Fig. 2a & 2b**). In PI440215, the STR were phased on both haplotypes of chromosome 5 (**Supplementary Fig. 10**.

### Gene prediction of assemblies and haplotypes

Gene prediction identified 69,601 genes (81,860 transcripts) in PI440215; and 72,890 genes (81,518 transcripts) in PGR7039. The average gene length in PI440215 is 3,084 bp, with the longest gene being 168,766 bp. In the case of PGR7039, the average gene length is 2,965 with the longest gene being 176, 255 bp. The BUSCO (Simão *et al*., 2015) score for these annotations was 98.7% and 98.9% for PI440215 and PGR7039, respectively (**Supplementary Fig. 4**).

Likewise, in the phased assembly of PI440215, we predicted 53,888 and 53,103 gene models in haplotype 1 and haplotype 2, respectively. The BUSCO completeness scores for these phased assemblies were 96.8% and 95.5%, respectively. Gene expression across haplotypes of PI440215 genome was compared to assess potential expression bias. Among 33,628 orthologous gene pairs; 4,697 genes exhibited expression bias (logFC>2, *P-value*<0.05) in at least one tissue type (leaf and stem). Overall, genes in haplotypes 2 showed higher expression levels compared with those in haplotype 1. The genes showing biased expressed span broad range of functional protein classes, with a strong representation of enzyme involved in primary and secondary metabolism, including transferase, oxidoreductase, hydrolase, kinase, ligases and proteases (**Supplementary Table 7**).

### Comparative genome analysis and genome evolution of bromegrass

Syntenic analysis was performed among the genomes of *B. tectorum, B. riparius*, *B. squarrosus, B. sterilis,* and *Hordeum vulgare* (**Fig. 3a**). All *Bromus* chromosomes exhibited a highly conserved syntenic architecture, indicating strong-macro level structural conservation across the genus. The well-known reciprocal translocation between chromosome 2H and 5H in *H. vulgare* was consistently detected when compared with each *Bromus* species, suggesting that *Bromus* retained the ancestral Triticeae chromosome arrangements. In *B. sterilis,* a major inversion on chromosome 1 was identified when compared with other *Bromus* species. Additionally, conserved gene duplications were detected across the *Bromus* species, particularly on chromosome 1 and 3, and 4 and 5. These results might suggest the presences of the ancient “rho” whole-genome duplication event in a broad range of grass species (Zhang et al., 2024).

**Fig. 3:**
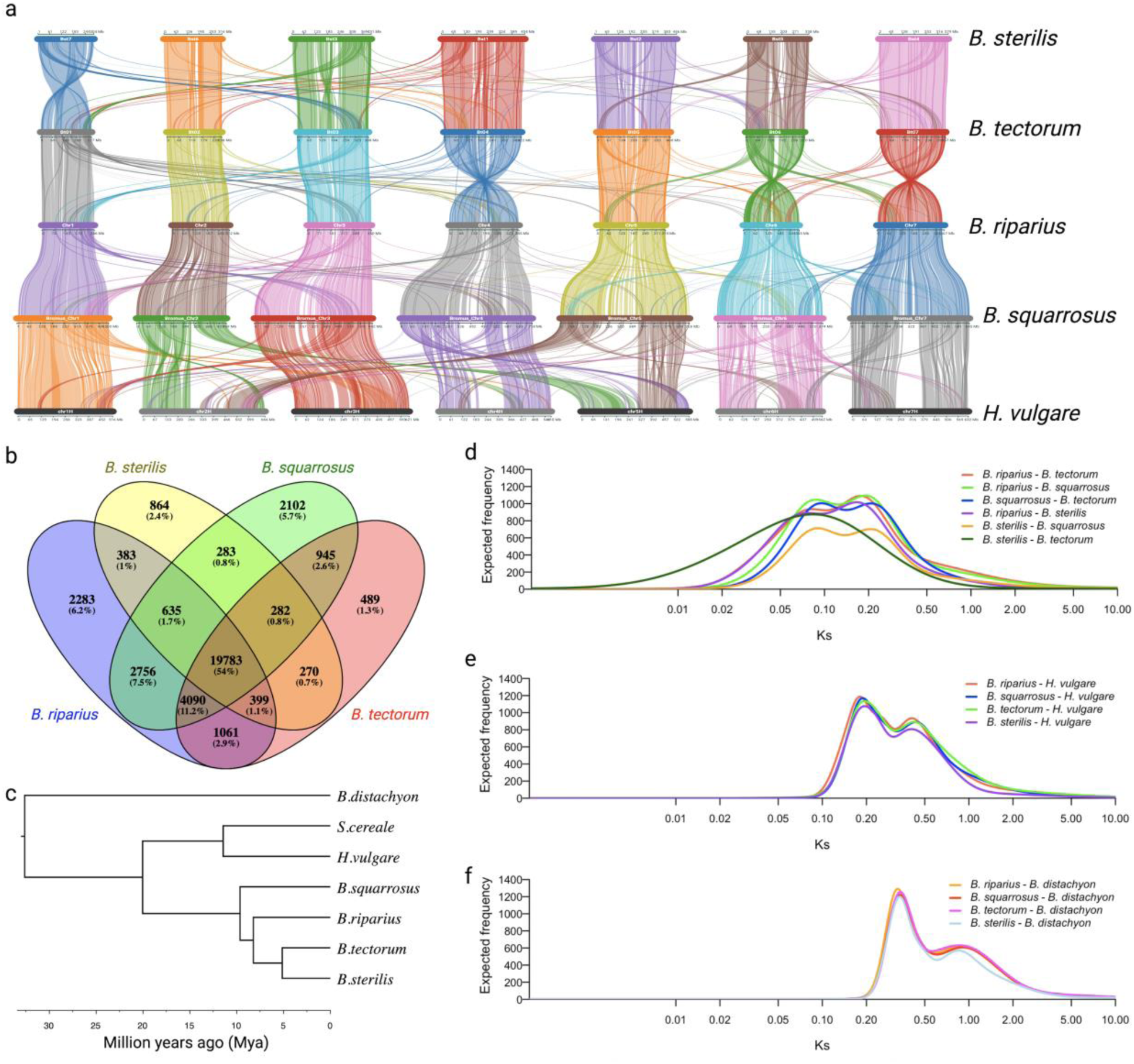
Comparative analysis of *Bromus* species. (a) Syntenic analysis among the *Bromus* species with *H. vulgare* showing an evolutionary conserved translocation on chromosome 2 and chromosome 5 and higher collinearity among the *Bromus* species. (b) Venn diagram showing core and dispensable orthogroups among *Bromus* species. (c) Species tree showing relationship among the *Bromus* species with other species under the Poaceae family. (d-f) Distribution of Ks to infer the evolution of genomes: (d) comparison among *Bromus* species, (e) comparison between *Bromus* species and *H. vulgare,* (f) comparisons between *Bromus* species and *B*. *distachyon*. Peaks indicate divergence events at distinct evolutionary time points.

Orthogroup analysis identified a total of 36,625 orthogroups in *Bromus* species (**Supplementary Table 8**). Among these, 19,783 represented core orthogroups; while 2,283, 489, 2,102 and 864 orthogroups were unique to *B. riparius*, *B. tectorum*, *B. squarrosus,* and *B. sterilis*, respectively (**Fig. 3b**). Overall, 58.74% of the orthogroups represented core groups, while 26.93% were classified as softcore and 14.32% as inter-dispensable orthogroups among the studied species (**Supplementary Table 9**). *Bromus tectorum* was found to have a higher loss of gene copies compared to all other *Bromus* species; *B. riparius* was found to have a higher gain in gene copies (**Supplementary Fig. 11**). A species tree was constructed based on single-copy orthologues and used to estimate divergence times through MCMCtree analysis (Reis and Yang, 2011). Further, Ks analysis of orthologs was performed to validate divergence estimates among the *Bromus* species. The analysis suggests that all *Bromus* species diverged from *Brachypodium distachyon* and *H. vulgare* approximately 33 and 20.27 Million years ago (Mya), respectively (**Fig. 3c**). Comparisons among the four diploid *Bromus* species revealed evidence of whole genome duplication events in all species except *B. sterilis* and *B. tectorum,* which appears to share the same lineage (**Fig 3d**). Within *Bromus*, *B. squarrosus* diverged earliest (9.75 Mya), followed by *B. riparius* (8.3 Mya), with *B. tectorum* and *B. sterilis* (5.2 Mya) diverging more recently. These relationships were further supported by the chloroplast-based phylogeny (**Supplementary Fig. 12**). Overall, all *Bromus* species exhibited conserved whole genome duplication patterns relative to *H. vulgare* and *B. distachyon* (**Fig. 3e and 3f**).

### Genetic diversity in bromegrass species

A total of 180 million SNVs were identified across 839 samples representing 18 different *Bromus* species. After filtering for sites with more than 20% missing data, a minor allele frequency below 1%, and heterozygosity greater than 90%, a final set of 14,395 high-quality SNVs was retained for downstream analysis. Population structure analysis identified six subpopulations within the genus *Bromus* (**Supplementary Table 10**). Distinct subpopulations was observed for lines representing *B. arvensis, B. squarrosus, B. racemosus*, as well as for double haploid (DH) lines derived from hybrid bromegrass. Additional subpopulations corresponded to *B. tectorum*, *B. hordeaceus, B. riparius*, and *B. inermis*. The hybrid population developed from meadow × smooth bromegrass displayed clear admixture from both parental types, whereas the DH lines formed a separate sub-population, likely reflecting reduced heterozygosity compared to other accessions (**Fig 4a**). Subpopulations representing *B. squarrosus, B. racemosus*, and *B. hordeaceous* were highly divergent, whereas cultivate *B. riparius* and *B. inermis* showed lower levels of genetic differentiation.

**Fig. 4:**
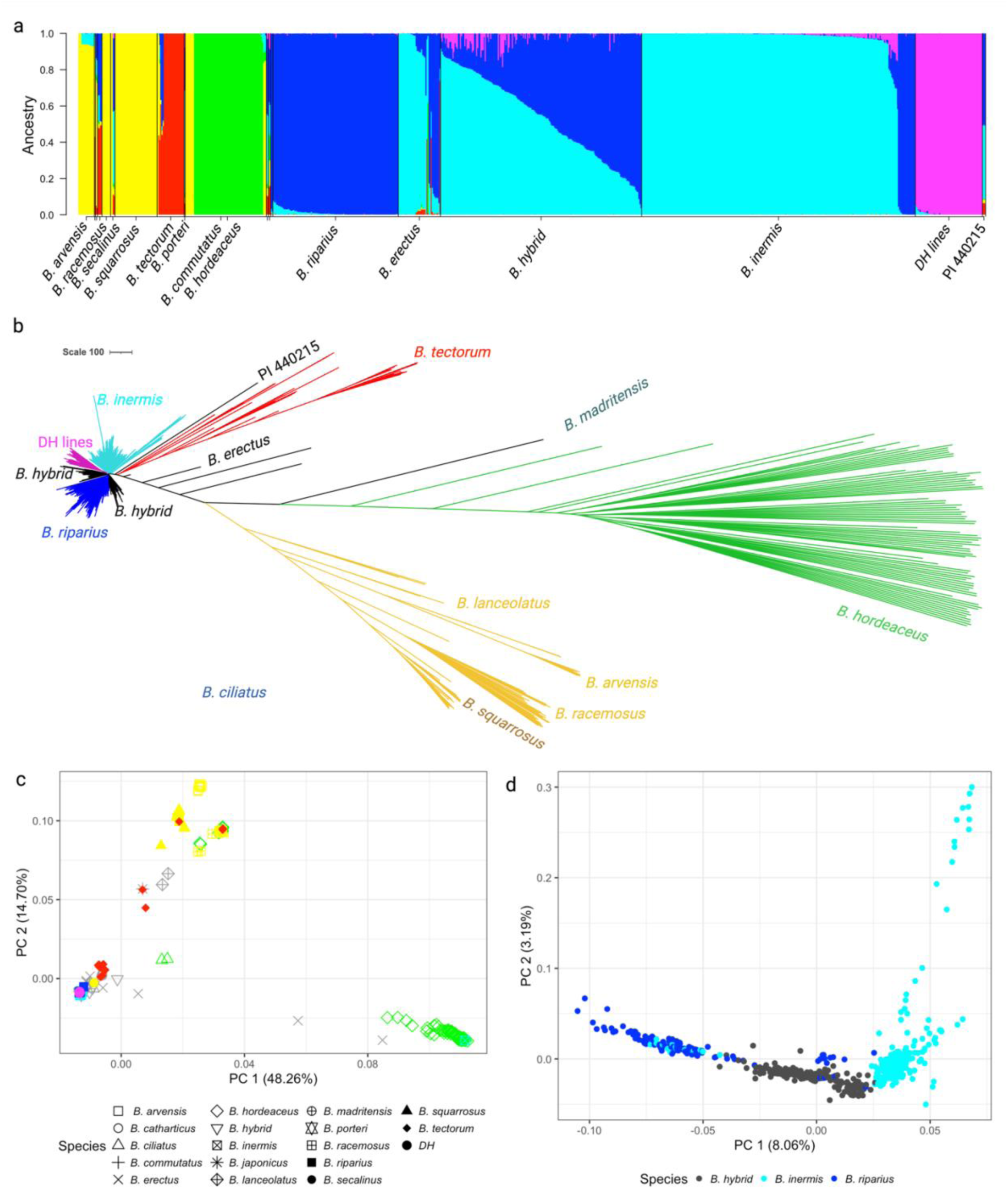
Genetic diversity in Bromegrass species. (a) Population structure analysis identified 6 populations, with two subpopulations corresponding to meadow bromegrass (blue) and smooth bromegrass (light blue). (b) Dendrogram shows genetic relationship among different bromegrass species, with higher divergence observed among wild relatives compared to cultivated bromegrass species. (c) Principal component analysis (PCA) showing 48.26% variation explained by the first component and 14.70% by the second component, the first component differentiates cultivated bromegrass from wild relatives. (d) PCA among cultivated bromegrass and breeding lines, differentiating hybrid bromegrass (black) developed from meadow (blue) and smooth bromegrass (light blue).

Phylogenetic analysis was performed using the Maximum Likelihood method with 1,000 replications in IQ-tree (Nguyen et al., 2015). The substitution model (GTR+F+ASC+R5) fitted as best model (Kalyaanamoorthy et al., 2017) to capture the effects of variable ploidy and high heterozygosity across samples. The resulting dendrogram showed a clear relationship among species and accessions, with a greater divergence among wild relatives compared to cultivated populations. Four major clusters were identified among the cultivated bromegrasses, representing *B. riparius*, *B. inermis*, *B. riparius* × *B. inermis* (hybrid) and DH lines. Further, wild relatives formed independent clusters reflecting the higher genetic distance among species (**Fig 4b**).

Principal Coordinate Analysis (PCoA) explained 48.26% of the variance in the first component and 14.70% in the second component, suggesting the high genetic diversity among the *Bromus* species (**Fig. 4c**). Cultivated bromegrass accession clustered together regardless of ploidy level. Further, a focused PCoA of *B. riparius*, *B. inermis*, *B. riparius* × *B. inermis* (hybrid) showed moderate levels of variation, with 8.06% and 3.19% explained by the first and second components, respectively (**Fig. 4d**). Breeding populations developed from *B. riparius* × *B. inermis* crosses displayed intermediate position consistent with admixture. Sub-population differentiation between *B. riparius* × *B. inermis* (hybrid) and two parents - *B. riparius* (F_ST_= 0.033), and *B. inermis* (F_ST_= 0.01) was low (**Supplementary Table 11**). Gene flow analysis further supported these patterns, revealing greater allele sharing between *B. inermis* to hybrid bromegrass than *B. riparius* and hybrid bromegrass (**Supplementary Fig. 13**).

## Discussion

Cultivated *Bromus* species are an important forage grass in North America for pasture and hay production, yet their large genome size, extensive ploidy levels, and poorly resolved subgenome structure make genomic studies challenging. Previous work relied on cytogenetic methods such as karyotyping (Klos et al., 2009; Tuna *et al*., 2001a) and flow cytometry to estimate genome in *Bromus* species. The recent availability of genome assemblies, including *B. tectorum* (Revolinski *et al*., 2023) and *B. sterilis* (Christenhusz *et al*., 2024), now enables comparative genomic investigation across the *Bromus* section. Genome size variation within *Bromus* is striking. Based on the results presented here the diploid *B. squarrosus* (PGR7039) has a genome size nearly twice that of diploid *B. riparius* (PI440215) (∼2.5 Gb), despite both species possessing the same chromosome number (Klos *et al*., 2009; Tuna et al., 2001b) with the higher proportion of repeat elements, particularly LTR elements (Angela, SIRE, Athila, CRM), driving the genome expansion in *B. squarrosus*. The centromere specific CRM element was one of the core repeat elements showing differences in PI440215 compared with PGR7039, suggesting that centromeric repeats play an important role in genome expansion dynamics. Understanding centromere organization is critical, as these regions are often underrepresented in genome assemblies but are essential for chromosome segregation and faithful recombination. This study supports a model in which centromere size scales with overall genome size in *Bromus,* highlighting their dynamic contribution to genome evolution (Chen et al., 2024; Plačková et al., 2024).

A particular notable finding in *B. riparius* (PI440215) is the presence of a unique high-order satellite structure consisting of a 390 bp monomer encapsulated within a 377 bp monomer, forming a distinct constriction-like structure. Such large STRs structure of this size have rarely been resolved in previous plant genome assemblies, as sub-telomeric regions are often collapsed or underrepresented due to their repetitive nature. Indeed, at a size of 27.7Mb the sequenced STR on chromosome 7 in PI440215 would surpass the 27.6Mb human satellite 3 (HSat3) repeat identified on chromosome 9 in the telomere-to-telomere human genome assembly that represents the largest such structure sequenced to date (Altemose et al., 2022). Although the functional significance of these structures remains unclear, their strong structural conservation could suggest a potential role in maintaining chromosomal stability. The intriguing possibility that the HSat3 array also has a functional role in gene expression by acting as a transcription factor binding platform for different pathways would provide another pivotal role for these structures (Franklin et al., 2025). In contrast, although similar satellite repeat motifs were identified in *B. squarrosus*, no comparable very large structural arrays were detected in the assembly, indicating lineage specific evolution of sub-telomeric repeats within the genus. The distinctive size and organization of these satellite structures also provide valuable markers that can be leveraged to differentiate subgenome composition in higher ploidy *Bromus* species.

Comparative repeat analysis revealed that Athila, Angela, and CRM elements have proliferated recently in *B. riparius*, but a higher abundance of these elements in *B. squarrosus* suggests independent evolutionary trajectory in these genomes. Despite these differences in repeats evolution, synteny remains high among *Bromus* species. An ancient translocation event involving chromosome 2 and 5 appears conserved across *Bromus* species compared to barley, likely occurring during divergence of *Bromus* from the genus *Hordeum.* Ks analysis supports a scenario in which *Bromus* lineage diverged around the same time, followed by independent evolutionary dynamics driving repeat expansion and speciation. However, the pangenome analysis revealed that *Bromus* species retained around 58% of their orthologs as core gene families, which is significantly higher than in closely related species such as *H. vulgare* and *B. distachyon* (Gordon et al., 2017; Jayakodi et al., 2024). This suggests that 1) despite dramatic differences in the repeat content the gene space remains relatively conserved with a high level of synteny maintained across the species, and 2) *B. squarrosus* diverged earlier to form a separate lineage possessing a higher proportion of repeat elements compared to *B. riparius* and *B. tectorum*. Allele-specific expression bias is common in plants, particularly heterozygous species, as reported in pears (Sun et al., 2025) and *Ficus carica* (Usai et al., 2025). Consistent with these observations, this study also identified widespread expression bias between alleles. The biased genes were predominantly associated with primary and secondary metabolic processes, which might suggest these imbalances may play an important role in regulating metabolic flexibility and adaptive responses in the genome.

Besides these foundational observations, the assembled genomes provide immediate utility for bromegrass breeding applications such as differentiation of germplasms at both the species level and at finer scales within breeding lines and utilization in the trait discovery. The identification of six subpopulation structures in *Bromus* species correlates with the taxonomic sections of the genus *Bromus*; however, the pronounced genetic differentiation between *B. inermis* and *B. riparius* suggests a historical divergence among the cultivated bromegrass species, although they are cross compatible (Coulman, 2004). This is particularly relevance for hybrid bromegrass, where distinguishing the genomic contribution of *B. riparius* and *B. inermis*, is critical for new variety development. The ability to identify admixtures and quantify subgenome contributions provides breeders with tools for marker assisted selection and genomic prediction. However, the absence of high-quality high ploidy genome assemblies continues to limit our understanding of subgenome interaction, homoeologous exchanges and dosage effects on gene expression. Sequencing of additional diploid lines and expanding to polyploid assemblies will be essential to capture the full spectrum genetic diversity in the genus *Bromus*.

## Methods

### Collection of germplasm

A total of 84 accessions were obtained from the Plant Gene Resources Center (PGRC), Saskatoon, along with 9 commercial cultivars, 62 DH lines developed at the National Research Council Canada (NRC), Saskatoon, 51 breeding lines from the Crop Development Center, University of Saskatchewan, and one diploid line from United States Department of Agriculture (USDA) genebank (**Supplementary Table 10**).

### DNA extraction and sequencing of diploid genomes

High Molecular Weight (HMW) DNA was extracted from single plants of diploid *B riparius* accession (PI440215) and diploid *B. squarrosus* accession (PGR7039) by using the NucleoBond® HMW DNA extraction kit (MACHEREY-NAGEL^TM^) for whole genome sequencing. For the remaining samples, genomic DNA was extracted using NucleoMag 96 kit (MACHEREY-NAGEL^TM^) on a KingFisher platform for genotyping. All the DNA samples were quantified using picogreen on a Tecan plate reader (Tecan Trading AG, Switzerland).

The diploid line PI440215 was sequenced using the long-read PacBio Sequel II platform (PacBio, Menlo Park, CA, USA) with five SMRT cells, while PGR7039 was sequenced using one SMRT cell on the Revio platform (PacBio, Menlo Park, CA, USA) in the Omics and Precision Analytics Laboratory (OPAL), GIFS, Saskatoon, Canada. Additionally, all samples were genotyped using a Genotyping by Sequencing (GBS) approach (Poland et al., 2012) and sequenced on a NovaSeq 6000 platform (Illumina, San Diego, CA, USA) in the OPAL.

### Hi-C library preparation and sequencing

Three Hi-C libraries for PI440215 and one Hi-C library for PGR7039 were prepared using the Dovetail® Omni-C^®^ kit and sequenced on the NovaSeq6000 platform. For the library preparation, approximately two grams of leaf samples were ground with the help of liquid Nitrogen, followed by crosslinking performed as recommended by the vendor provided protocol (Cantata Bio, Scotts Valley, CA, USA). The quality of library was assessed by using the BioAnalyzer High Sensitivity Kit (Agilent Technologies, Santa Clara, California, USA)

### Whole genome assembly and scaffolding

The raw HiFi sequence data obtained from PacBio were assembled using hifiasm 0.19.5 (Cheng *et al*., 2022) with default parameters except n = 9. The resulting base assemblies were scaffolded using Hi-C data generated with the Omni-C protocol. HiC-Pro (Servant et al., 2015) was employed to map the Hi-C reads to the base assembly, and EndHiC (Wang et al., 2022) was used for scaffolding the contigs in chromosome level. The 3D-DNA (Dudchenko et al., 2017) was utilized to generate Hi-C files, which were visualized in Juicebox (Durand et al., 2016). Manual curation was performed in Juicebox as needed.

### Illumina library preparation and sequence analysis

DNA libraries for PI440215 and PGR7039 were prepared using the PCR free DNA Library Preparation Kit (Illumina, San Diego, CA, USA) and sequenced on a NovaSeq600 platform. The obtained reads were trimmed using Trimmomatic v.0.39 (Bolger et al., 2014) for low quality and reads shorter than 51 bp, retaining paired-end reads only. The Illumina reads were utilized to calculate the K-mer frequencies using Jellyfish (Marçais and Kingsford, 2011), and the genome sizes were estimated using GenomeScope (Vurture *et al*., 2017). The Illumina short reads were mapped to the respective assembled genome using BWA v.0.7.18 (Li and Durbin, 2009) and mapping quality was assessed using Qualimap (García-Alcalde et al., 2012). The read distributions were plotted against the assembled genome using circos (Krzywinski et al., 2009).

### RNA Extraction, library construction, sequencing, and analysis

RNA extraction was performed from two tissue types (stem and leaves) for both PI440215 and PGR7039. The three biological replicates for each tissue type were extracted using RNeasy Plant Qiagen kit (Qiagen) according to the manufacturer’s protocol. On-column DNA digestion was performed for all samples, and the quantity and quality of total RNA were assessed using the BioAnalyzer RNA kit (Agilant Technologies, Santa Clara, California, USA). RNA libraries were prepared by using the Illumina Stranded mRNA Prep kit (Illumina, San Diego, CA, USA) with an initial RNA input of 500 ng per sample. The RNA libraries were quantified using the BioAnalyzer High Sensitivity Kit (Agilant Technologies, Santa Clara, California, USA) and sequenced on the NovaSeq 6000 platform. The same RNA samples were also sequenced on the PacBio Revio platform using the Kinnex Kit (PacBio, Menlo Park, CA, USA) to obtain full-length transcripts. All the RNA-seq reads were trimmed for low-quality and reads shorter than 51 bp using Trimmomatic v.0.39 (Bolger *et al*., 2014), retaining only paired-end reads. The filtered reads were mapped to their respective genomes using STAR v. 2.7.9a (Dobin et al., 2013), and transcript abundance was quantified using RSEM v.1.3.1 (Li and Dewey, 2011). Differential gene expression between haplotypes was assessed using a *t*-test across three biological replicates following ortholog identifying. Orthologs of differentially expressed genes were identified against the *Hordeum vulgare* cv. Morex V3 (Mascher et al., 2021) and subjected to functional classification the PANTHER online tool (Thomas et al., 2022).

### Gene and repeat annotation

The machine learning based helixer tool (Holst et al., 2023) and *de novo* Braker v.2.1.6 (Brůna et al., 2021) were used to predict gene and gene structure. In the case of Braker annotation protein evidence from *Arabidopsis thaliana*, *Brachypodium distachyon*, *Hordeum vulgare*, *Secale cereal*, *Oryza sativa* were used along with the evidence from RNA seq data generated from the leaves and stem tissues. Further, Kinnex data were mapped to the genome using pbmm2 v.1.14.0 after collapsing the reads with isoseq v.4.1.2. The resulting gene structure information from helixer, Braker v.2.1.6, and Kinnex data was merged to create the final version of gene annotation. Gene annotation completeness was evaluated using BUSCO v.5.2.2 (Simão *et al*., 2015) with embryophyte_odb10 database.

Repeat analysis was conducted using the *de-novo* based TE annotation tool EDTA v.2.1.0 (Ou *et al*., 2019), and the raw TE library was developed. The raw TE libraries from *B. tectorum, B. riparius*, *B. squarrosus, B. sterilis* were concatenated and curated for the unknown and similar repeats present across the *Bromus* species. The final repeat library consists of 27,122 repeat elements which were used to mask the genome using RepeatMasker v.4.1.2-p1 (Tarailo-Graovac and Chen, 2009). LTR_retriever which is part of EDTA pipeline was employed to identify full-length LTR, and TEsorter (Zhang et al., 2019) was used to classify the full-length LTR in to specific clades. LAI was employed to calculate the LTR Assembly Index (Ou *et al*., 2018). Further, tandem duplicate genes were predicted using the K-mer based repeat annotation tool, TRASH (Wlodzimierz et al., 2023), Manual classification of tandem repeats was performed based on the size, and their visualization was performed in R statistical software (Team, 2021) with a package karyoploteR (Gel and Serra, 2017). Further the tandem repeats of same class were aligned using software MEGA v.11 (Tamura et al., 2021) to develop a phylogenetic tree, which was visualized using iTOL v. 7.1 (Letunic and Bork, 2024).

### Orthogroup analysis

The annotated genes from PI440215 and PGR7039, along with published genome of *B. tectorum* (Revolinski *et al*., 2023), *B. sterilis* (Christenhusz *et al*., 2024)*, H. vulgare* (Mascher *et al*., 2021)*, Secale cereal* (Rabanus-Wallace et al., 2021) and *B. distachyon* (International Brachypodium, 2010), were used to identify orthogroups using OrthoFinder v.2.5.4 (Emms and Kelly, 2019). Single-copy orthogroups were utilized to construct a species tree to determine the genetic relationship among the species. Further, gene family loss and gains were analysed using CAFE v.5.2.2 (Mendes et al., 2020). MCMC tree (Reis and Yang, 2011) with a 2000 burn-in, and a calibration point of 29.3 -35.5 Mya for *B. distachyon* and *Bromus* and 6.8 -18.3 mya between *H. vulgare* and *S. cereal* were used for identifying age of divergence among species.

### Synteny analysis

Synteny analysis was performed with annotated genes of diploid bromegrass PI440215 and PGR7039 along with the published genomes of *Bromus sterilis, Bromus tectorum*, *Hordeum vulgare* genomes. First, a BLASTp all-versus-all comparison was performed for the predicted protein for all species and the protein blast results were utilized generate a syntenic file in MCScanX (Wang et al., 2012) with default parameters and results were visualized using an online platform SynVisio (Bandi and Gutwin, 2020).

### Synonymous substitution analysis

Gene pairs identified from the syntenic analysis were further analysed for synonymous substitution rates using the GenoDup pipeline (Mao, 2019); where protein sequences were align using MAFFT (Katoh et al., 2002), and Ks value were calculated using codeml package in PAML (Yang, 2007). A gaussian mixture model was implemented in R/mclust (Scrucca et al., 2016) to fit the distributions.

### Chloroplast assembly and phylogenetic analysis

The long-read were used to assembled the chloroplast using Oatk (Zhou et al., 2024). (**Supplementary Table 12**). The chloroplast genome maps were constructed using the OGDRAW program (Greiner et al., 2019). The assembled chloroplast genomes were aligned with published chloroplast genome of *Brachypodium distachyon, Triticum aestivum, Thinopyrum elongatum, Hordeum vulgare*, *Bromus vulgaris*, and *Oryza sativa* using clustal omega (Sievers et al., 2011). Maximum likelihood tree construction and visualization was performed in MEGA11 (Tamura *et al*., 2021).

### GBS data analysis

Raw reads were demultiplexed using an in-house script. Demultiplexed samples were trimmed for low-quality reads using Trimmomatic v.0.39 (Bolger *et al*., 2014), with reads shorter than 51 bp removed. The reads were mapped to the *Bromus riparius* genome (PI440215) using BWA v.0.7.18 (Li and Durbin, 2009). Variants were called using BCFtools (Danecek et al., 2021).

Population structure was analysed using STRUCTURE software v.2.3.4 (Pritchard et al., 2000) with three replications for each K=1 to K=10 with MCMC of 50,000 and burn-in of 50,000, while principal component analysis was conducted using TASSEL v.5.0 (Bradbury et al., 2007). The structure harvester was used to determine the number of subpopulations using Evanno method (Earl and VonHoldt, 2012; Evanno et al., 2005) (**Supplementary Table 13**). A phylogenetic tree was constructed using Maximum Likelihood method with 1,000 replications in IQ-tree v2.4.0 (Minh et al., 2020), to infer genetic relationships among the studied lines. Sub-population differentiation was calculated using vcftools (Danecek et al., 2011).

To identify admixed taxa in Bromegrass populations (PI440215, *B. inermis*, *B. erectus*, *B. hybrid*, *B. riparius*), five accessions including PI440215, CN32686, CN32371, S9629HB_3-16, and Fleet were selected based on the high Q value representing each species, and CN32344 (*B. ciliatus*) was used as an outgroup. The filtered VCF dataset including six selected samples was converted to fasta format by using vcf2phylip (Ortiz, 2019). IQ-tree v2.4.0 was used to construct a species-tree using maximum-likelihood method with bootstrapping of 1000 replications (Minh et al., 2020). The inferenced species-tree was used as a proposed tree for Dsuite program. The gene flow analysis was performed with Dsuite software (Malinsky et al., 2021). Dtrios command was used to calculate the D statistics and the f4-ratio for all combinations of trios of species. The f-branch metric was calculated with subcommand Fbranch indicating the gene flow between specific branches. f-branch values were plotted using a python script (dtools.py) in the same package of Dsuite.

## Acknowledgements

We would like to thank the Agriculture Development Fund, Saskatchewan Ministry of Agriculture (ADF 20200420) and the Saskatchewan Forage Seed Development Commission for financial support. We would also like to thank Byambatseren Dashnyam for sampling bromegrass lines for genetic diversity studies.

## Data Availability

Raw sequencing data have been deposited in National Center for Biotechnology Information (NCBI) under the BioProject accession PRJNA1391830, and the supplemental materials are available on Figshare at 10.6084/m9.figshare.31008613.

## Supplementary Figures

Supplementary Fig. 1: Estimation of genome size using K-mer Analysis. A) PI440215 and B) PGR7039.

Supplementary Fig. 2: Flow cytometry-based estimation of genome size across *Bromus* species. Sample 2-2-5 corresponds to PI440215, and sample 7039 corresponds to PGR7039. Barley, wheat, and a higher ploidy bromegrass (CN1086781) were used as standards.

Supplementary Fig. 3: Contact map showing HiC scaffolding of PI440215 (*Bromus riparius*) haplotypes. A) Haplotype 1 and B) Haplotype 2.

Supplementary Fig. 4: BUSCO completeness analysis of bromegrass genomes assemblies and annotations for PI440215, its two haplotypes (haplotype 1 and haplotype 2), and PGR7039.

Supplementary Fig. 5: Distribution of full-length LTR in bromegrass genomes: A) PI440215 and B) PGR7030. Tracks from top to bottom represents CRM (red), SIRE (light blue), Retand (purple), Athila (green), Angela (grey), and Tekay (blue), centromeric positions are represented as a black band on the ideogram with corresponding physical positions.

Supplementary Fig.6: Comparison of proliferation age of Copia and Gypsy retrotransposon elements in PI440215 (blue) and PGR7039 (red). Multiple peaks indicate proliferation events at different time points for the Angela (d), Athila (e), Retand (f), SIRE (g), CRM (h) and Tekay (i) elements.

Supplementary Fig. 7: Distribution of telomere, centromere specific CRM, and NOR (45S rDNA) repeats in PI440215 (a) and PGR7039 (b).

Supplementary Fig. 8: Confirmation of satellite repeat structures on chromosome 5 and 7 in *Bromus riparius* from Hi-C data. Green indicates contig boundaries, while purple indicates chromosome boundaries. The presence of single contigs overlapping satellite structure and the remainder of the chromosome suggests that these satellite repeats (STR) are integral components of the assembled chromosome.

Supplementary Fig. 9: Variability of the 390-mer (A) and 377-mer (B) tandem repeats associated with satellite structure in *Bromus riparius* across bromegrass species. Most tandem repeats variants were species specific (390-mer) as well as shared among groups (377-mer). Phylogenetic trees were constructed based on multiple sequence alignments of the 390-mer and 377-mer sequences identified across bromegrass species. The abundance of each sequence variant is shown as a bar plot, with the count values displayed at the tree tips.

Supplementary Fig. 10: Genome wide distribution of tandem repeats in PI440215 haplotype 1 (A) and haplotype 2 (B). Coloured bands represent tandem repeats of different length, as indicated in the legend.

Supplementary Fig 11. Estimation of gene family loss and gain events across Bromus species. Gene family loss and gain events are summarized on the branches where positive figures represent gain and negative figures represent loss.

Supplementary Fig. 12: A) Circular map of the *Bromus riparius* chloroplast genome showing the distribution of protein coding genes, rRNA, and tRNA. Functional groups are color coded according to the legend. The innermost circle represents the relative GC (dark) and AT (light) content of the chloroplast genome. B) Phylogenetic relationships among *Bromus* species and related taxa inferred from chloroplast genome sequences. Accession numbers are indicated at the nodes for publicly available chloroplast genome assemblies.

Supplementary Fig.13. Pattern of gene flow among cultivated bromegrass accessions inferred from GBS data.

## Supplementary Tables

Table S1. Assembly statistics of *Bromus riparius* (PI440215) and *B. squarrosus* (PGR7039).

Table S2. Mapping statistics of Illumina short-reads to the genome of *Bromus riparius* (PI440215) and *B. squarrosus* (PGR7039).

Tables S3. Classification and proportion of repeats elements in the genome of *Bromus riparius* (PI440215), haplotypes of PI440215, *B. squarrosus* (PGR7039), *B. tectorum*, and *B. sterilis*.

Table S4. Classification of Full-length LTR in the genome of *Bromus riparius* (PI440215) and *B. squarrosus* (PGR7039).

Table S5. Identification of tandem repeats in *Bromus riparius* (PI440215).

Table S6. Identification of tandem repeats in *Bromus squarrosus* (PGR7039).

Table S7. Gene expression biases in haplotypes of *Bromus riparius* (PI440215).

Table S8. Orthogroup analysis in *Bromus* species and comparison with *Secale cereal*, *Hordeum vulgare* and *Brachypodium distachyon*.

Table S9. Classification of orthogroups across *Bromus* species. Table S10. Q-matrix for six-subpopulations in *Bromus* species.

Table S11. Subpopulation differentiation among cultivated Bromegrass lines.

Table S12. Chloroplast assembly statistics of *Bromus riparius* (PI440215) and *B. squarrosus* (PGR7039).

Table S13. Determination of the number of subpopulations in *Bromus* species.

