## Supplemental Figures for "High-quality genome assemblies of diploid *Bromus* species enhance understanding of genome complexity and uncover large DNA satellite structures"

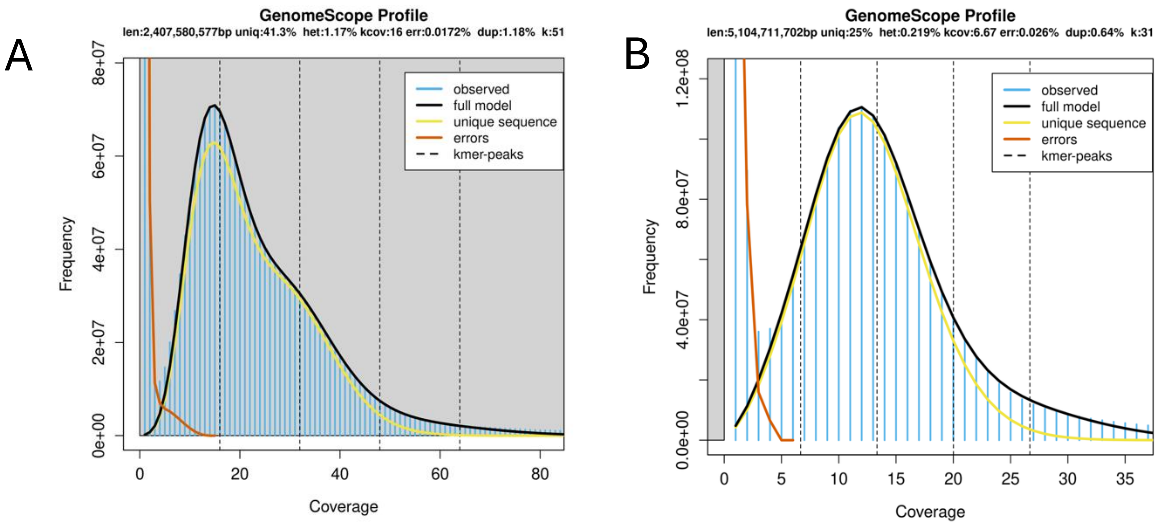


Supplementary Fig. 1: Estimation of genome size using K-mer Analysis. A) PI440215 and B) PGR7039.

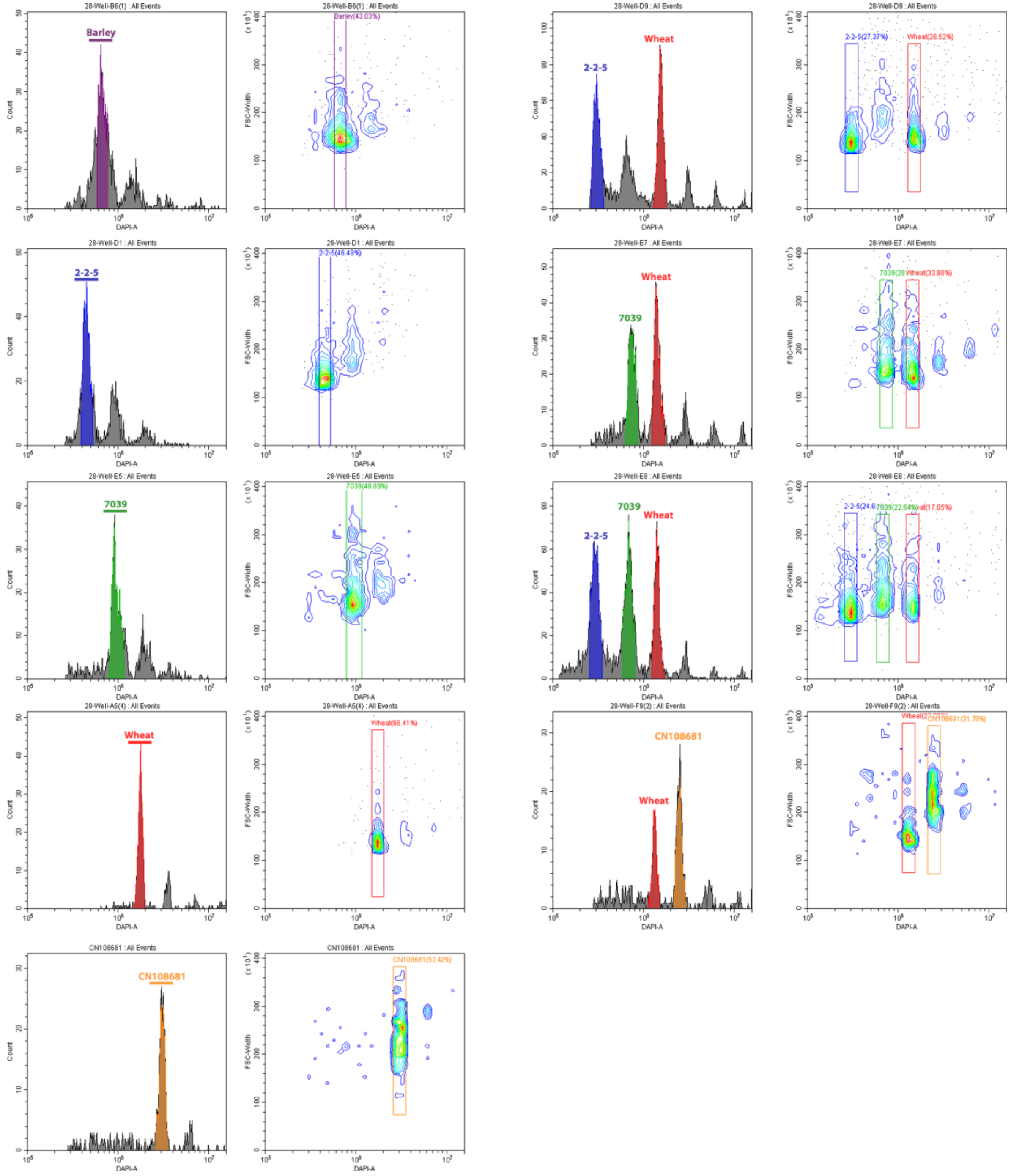


Supplementary Fig. 2: Flow cytometry-based estimation of genome size across *Bromus* species. Sample 2-2-5 corresponds to PI440215, and sample 7039 corresponds to PGR7039. Barley, wheat, and a higher ploidy bromegrass (CN1086781) were used as standards.

A B

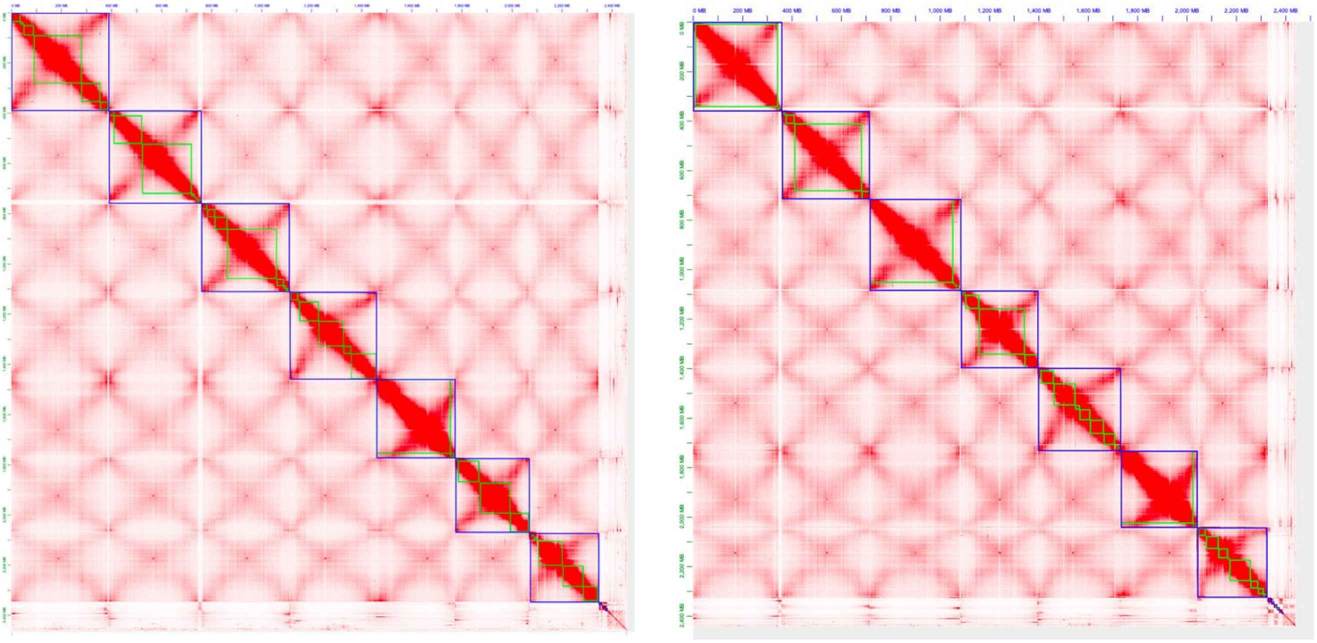

Supplementary Fig. 3: Contact map showing HiC scaffolding of PI440215 (*Bromus riparius*) haplotypes. A) Haplotype 1 and B) Haplotype 2.

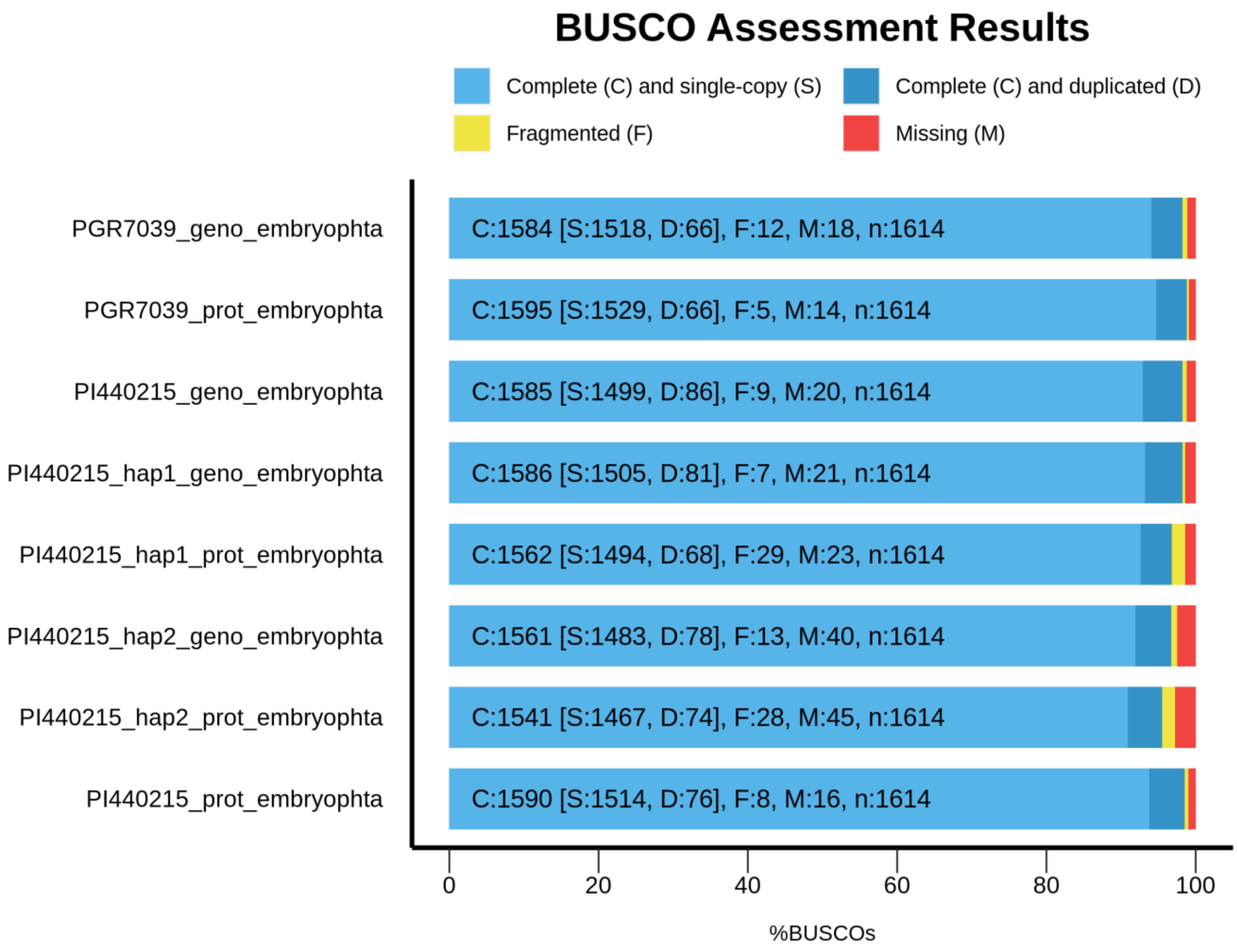


Supplementary Fig. 4: BUSCO completeness analysis of bromegrass genomes assemblies and annotations for PI440215, its two haplotypes (haplotype 1 and haplotype 2), and PGR7039.

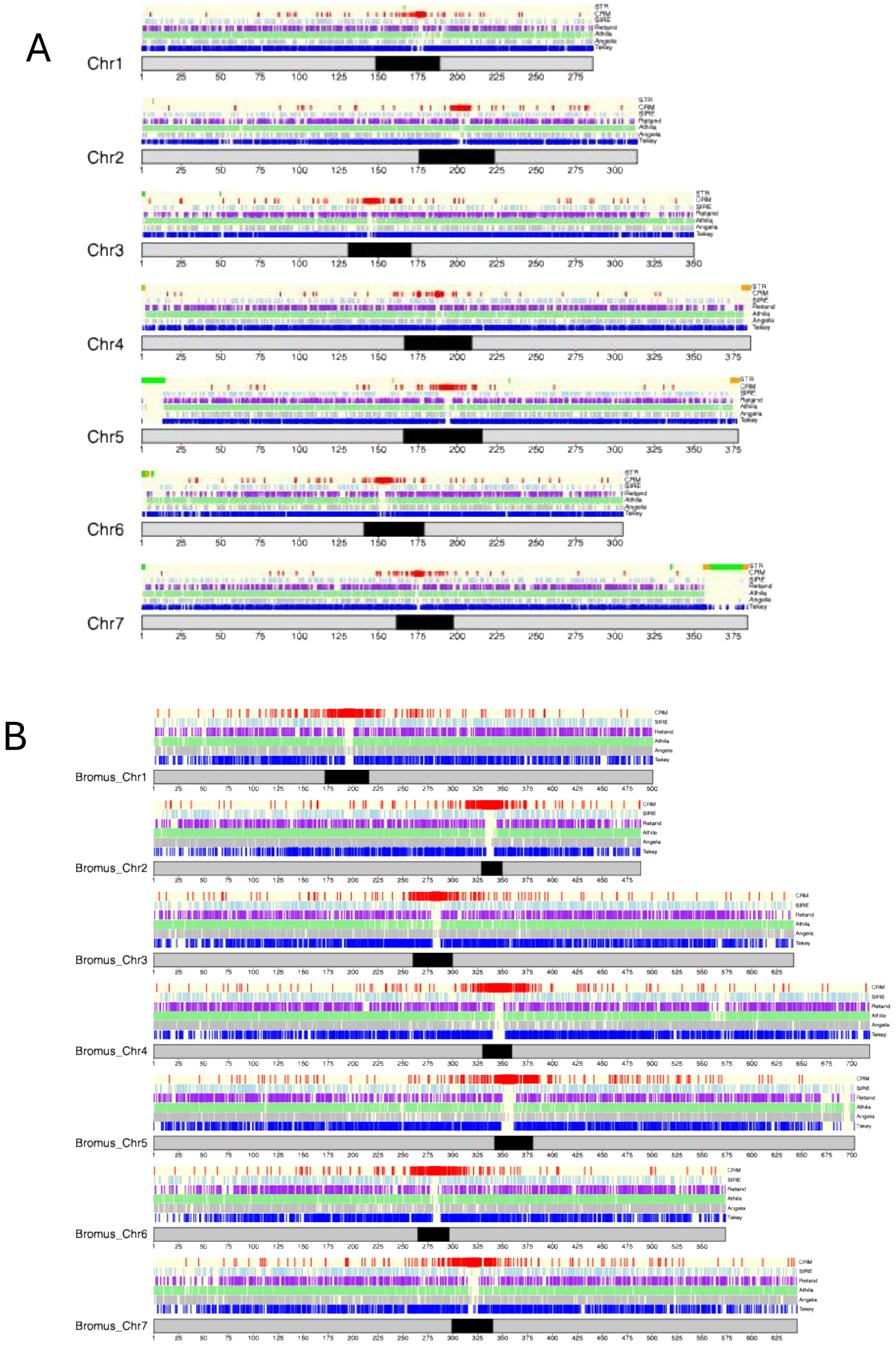


Supplementary Fig. 5: Distribution of full-length LTR in bromegrass genomes: A) PI440215 and B) PGR7030. Tracks from top to bottom represents CRM (red), SIRE (light blue), Retand (purple), Athila (green), Angela (grey), and Tekay (blue), centromeric positions are represented as a black band on the ideogram with corresponding physical positions.

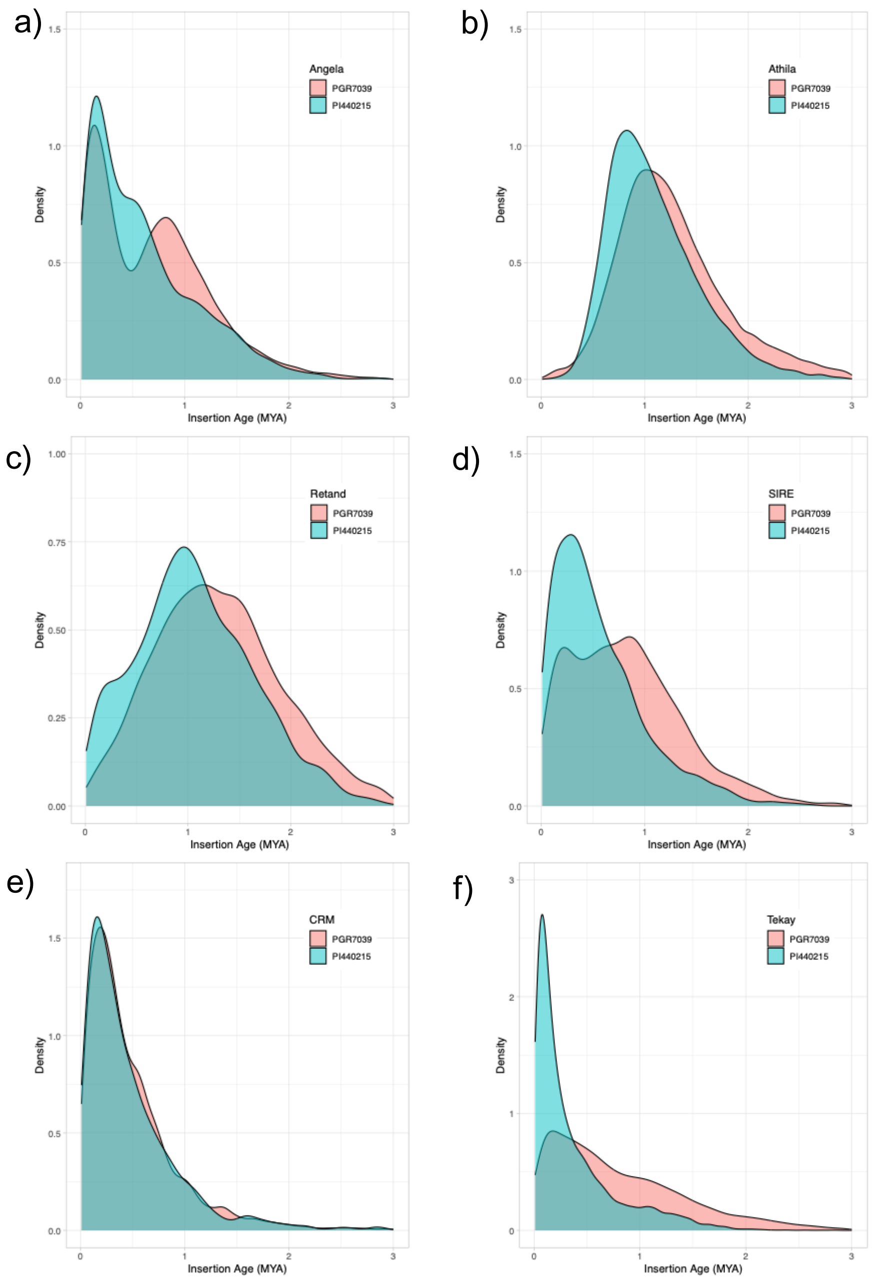


Supplementary Fig. 6: Comparison of proliferation age of Copia and Gypsy retrotransposon elements in PI440215 (blue) and PGR7039 (red). Multiple peaks indicate proliferation events at different time points for the Angela (d), Athila (e), Retand (f), SIRE (g), CRM (h) and Tekay (i) elements.

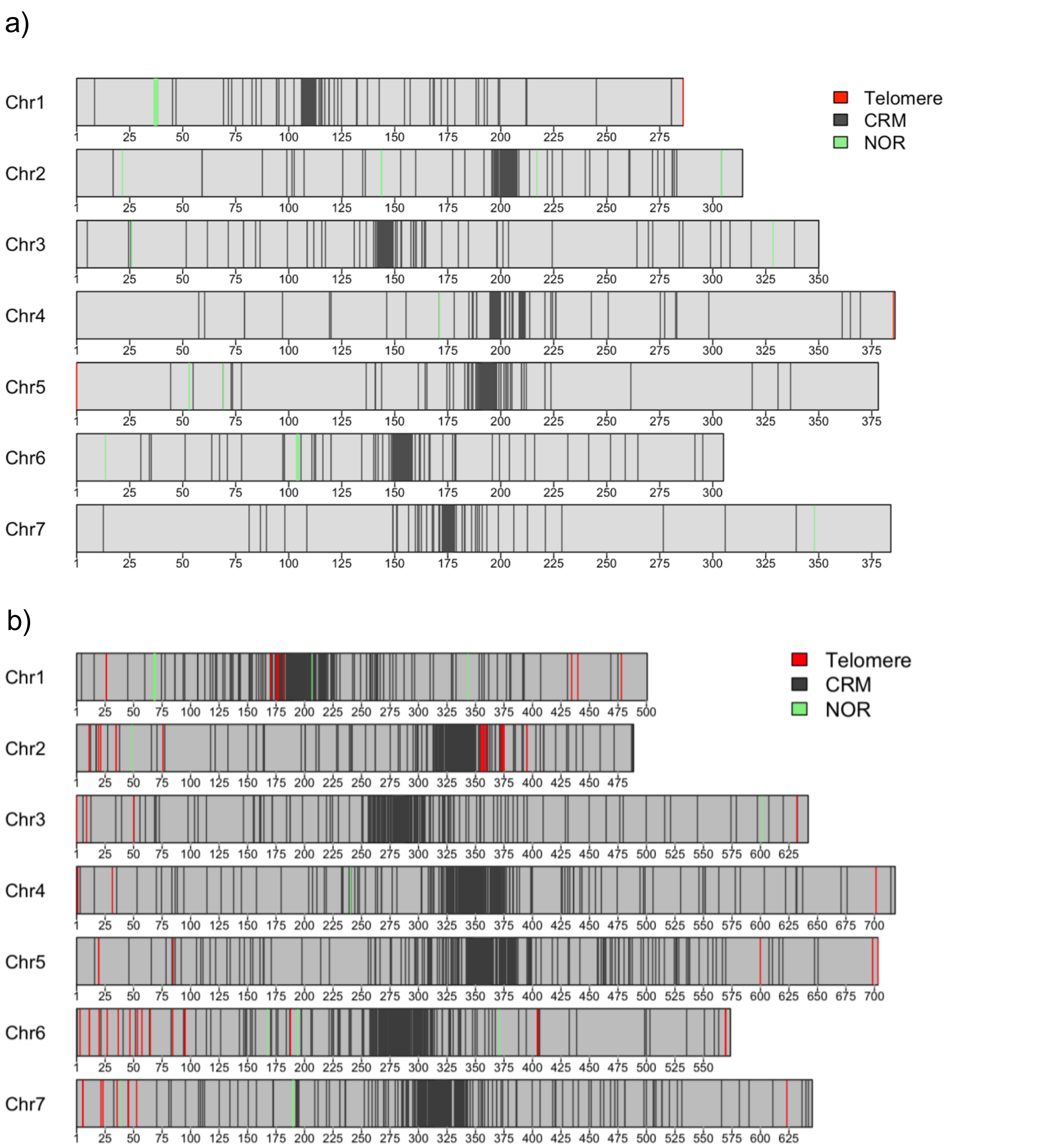


Supplementary Fig. 7: Distribution of telomere, centromere specific CRM, and NOR (45S rDNA) in PI440215 (a) and PGR7039 (b).

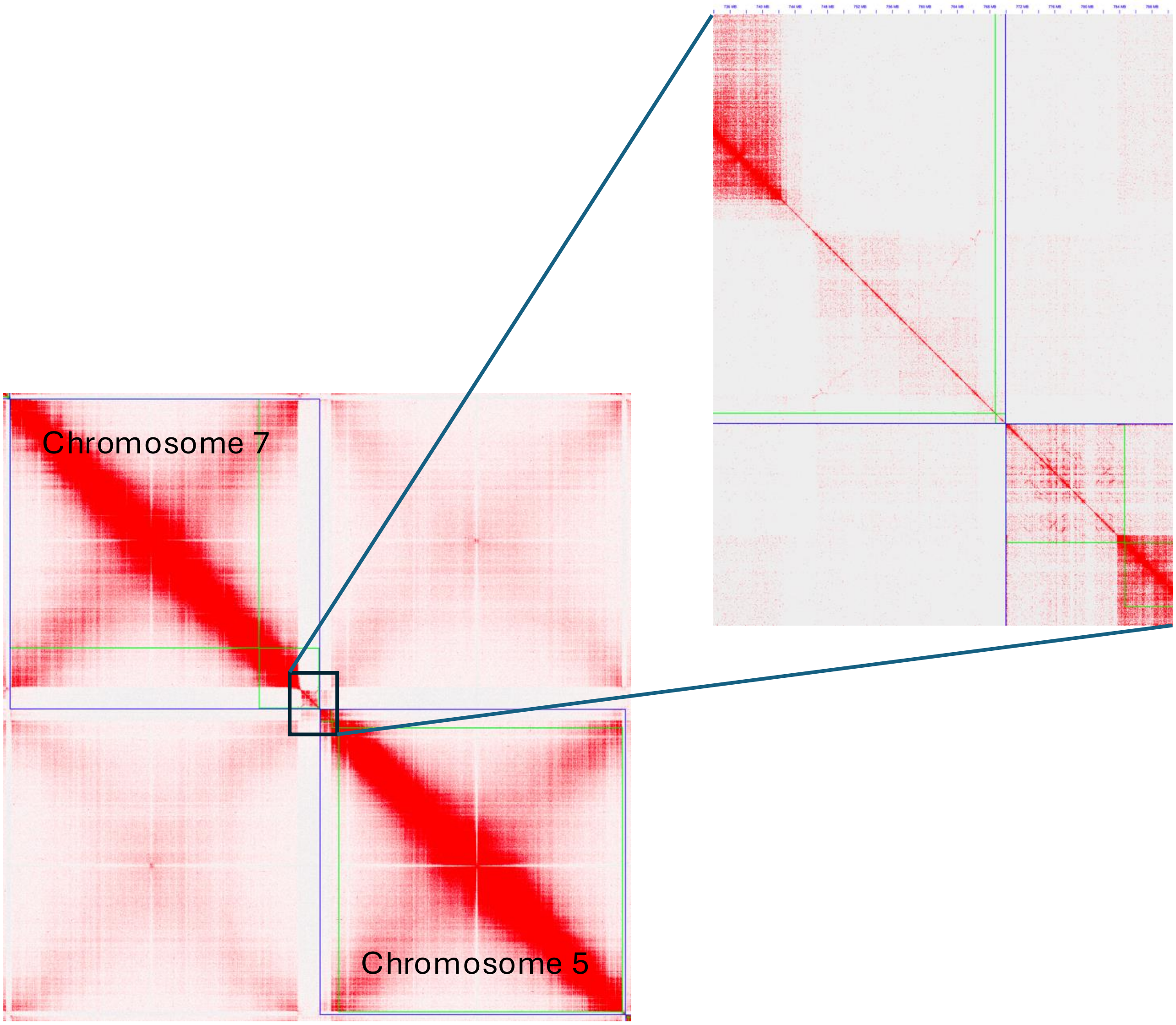


Supplementary Fig. 8: Confirmation of satellite repeat structures on chromosome 5 and 7 in *Bromus riparius* from Hi-C data. Green indicates contig boundaries, while purple indicates chromosome boundaries. The presence of single contigs overlapping satellite structure and the remainder of the chromosome suggests that these satellite repeats (STR) are integral components of the assembled chromosome.

A. B.

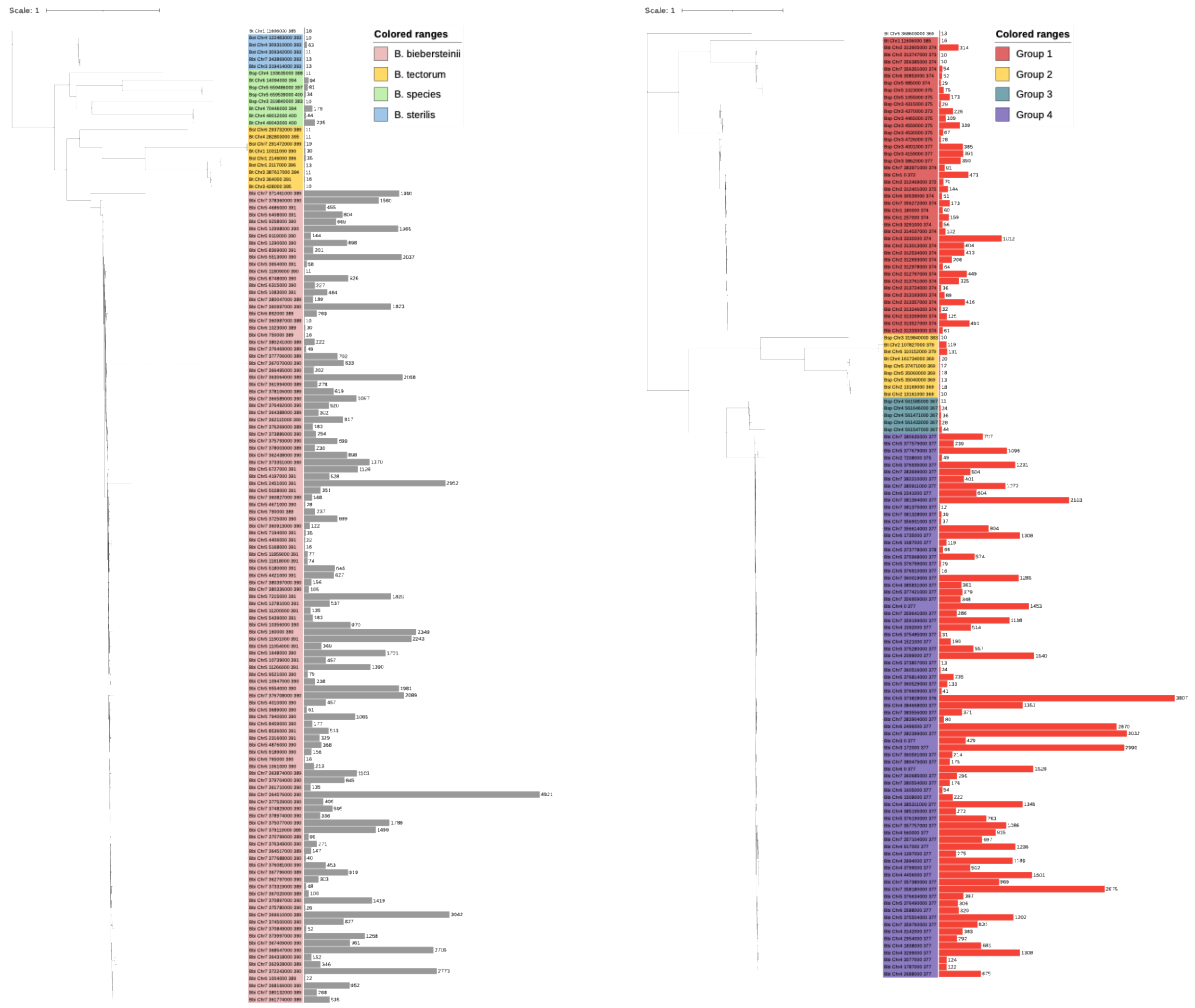


Supplementary Fig. 9: Variability of the 390-mer (A) and 377-mer (B) tandem repeats associated with satellite structure in *Bromus riparius* across bromegrass species. Most tandem repeats variants were species specific (390-mer) as well as shared among groups (377-mer). Phylogenetic trees were constructed based on multiple sequence alignments of the 390-mer and 377-mer sequences identified across bromegrass species. The abundance of each sequence variant is shown as a bar plot, with the count values displayed at the tree tips.

A

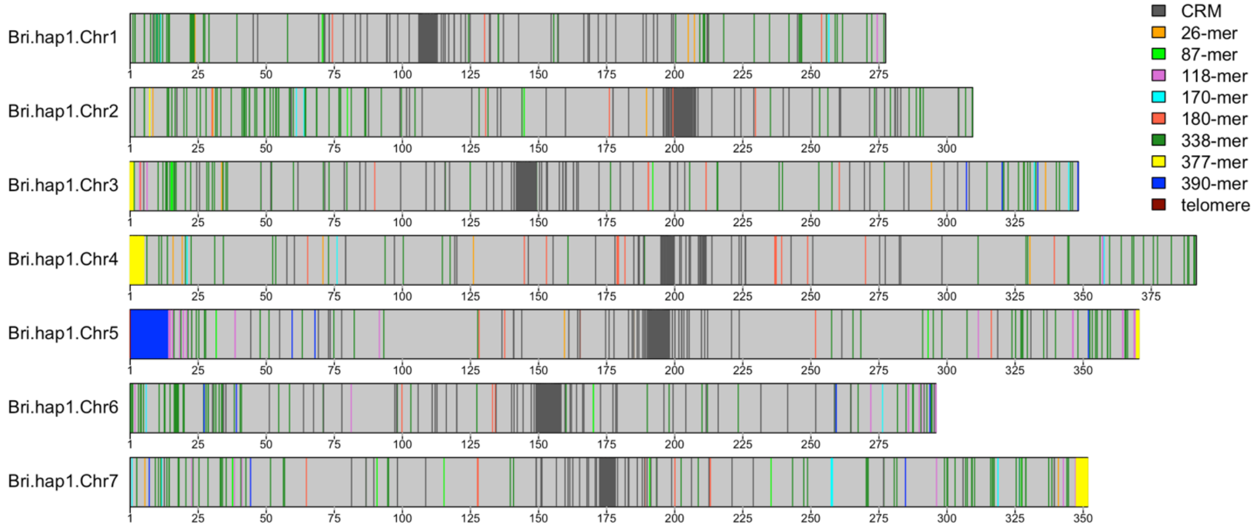


B

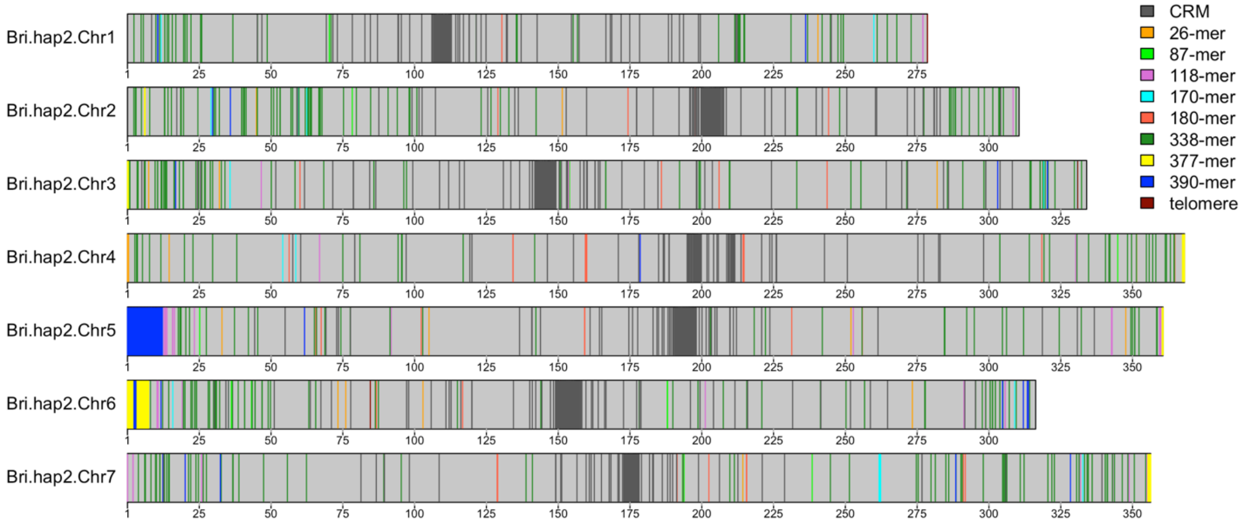


Supplementary Fig. 10: Genome wide distribution of tandem repeats in PI440215 haplotype 1 (A) and haplotype 2 (B). Coloured bands represent tandem repeats of different length, as indicated in the legend.

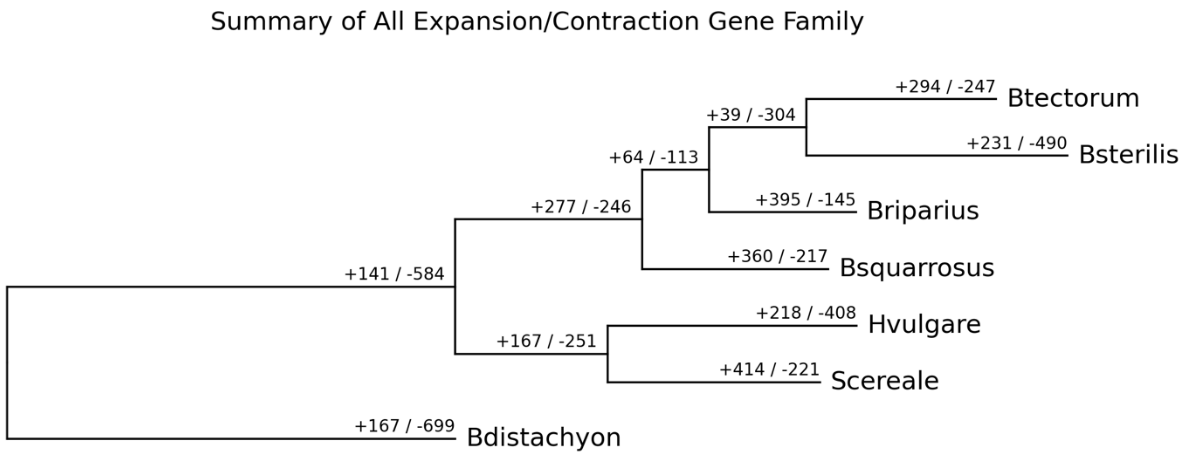


Supplementary Fig 11. Estimation of gene families loss and gain event across Bromus species. Gene families loss and gain events are summarized on the branches where positive figure represent gain and negative figure represent loss of number of gene families.

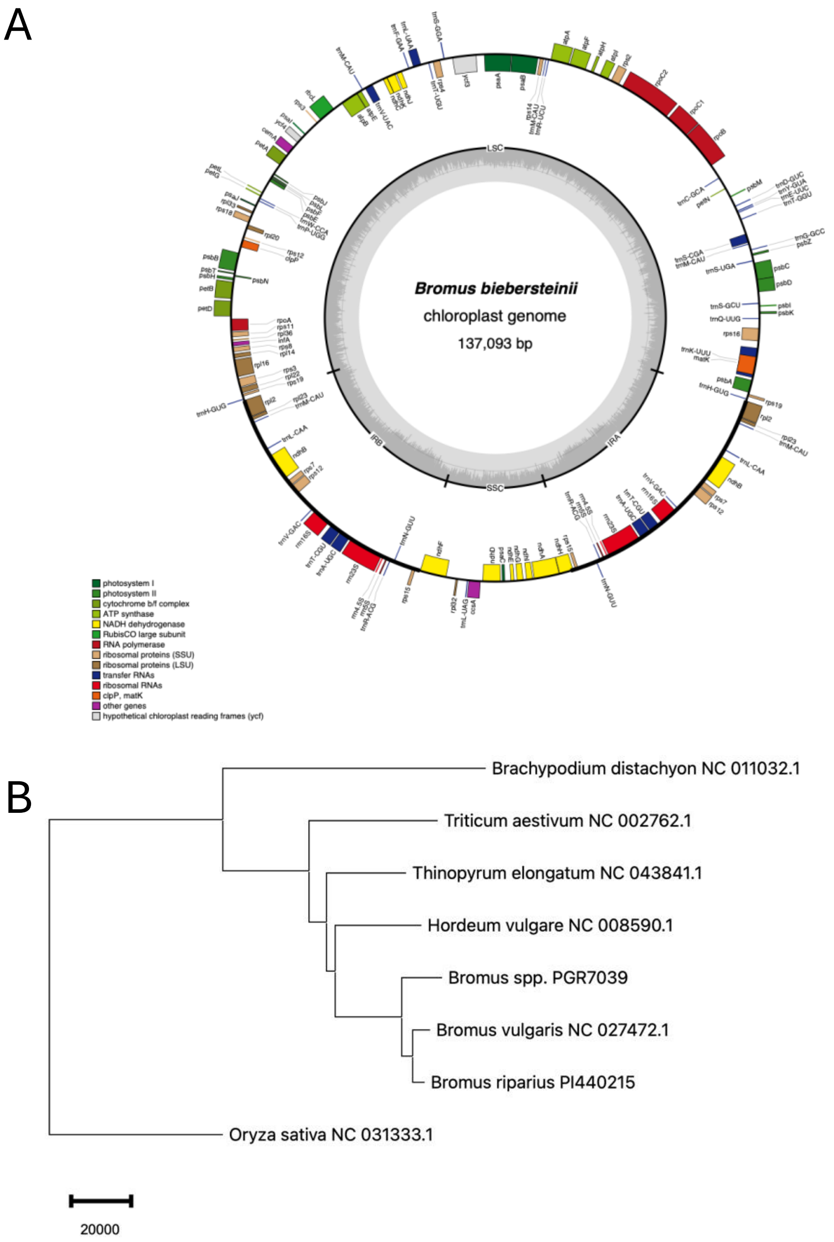


Supplementary Fig. 12: A) Circular map of the *Bromus riparius* chloroplast genome showing the distribution of protein coding genes, rRNA, and tRNA. Functional groups are color coded according to the legend. The innermost circle represents the relative GC (dark) and AT (light) content of the chloroplast genome. B) Phylogenetic relationships among *Bromus* species and related taxa inferred from chloroplast genome sequences. Accession numbers are indicated at the nodes for publicly available chloroplast genome assemblies.

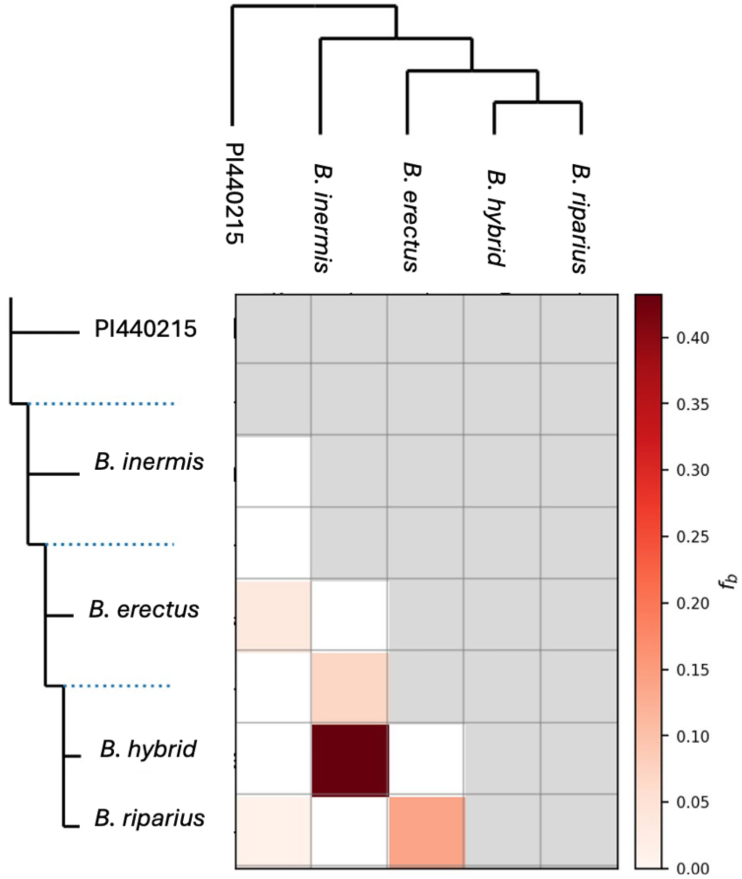


Supplementary Fig.13. Pattern of gene flow among cultivate bromegrass accession inferred from GBS data.
